# KIF13B controls ciliary protein content by promoting endocytic retrieval and suppressing release of large extracellular vesicles from cilia

**DOI:** 10.1101/2024.03.21.586066

**Authors:** Csenge K. Rezi, Alina Frei, Fabiola Campestre, Karsten Boldt, Benjamin Mary, Anna Maria Fixdahl, Anna-Louise With Petersen, Aurelien Sicot, Christina R. Berggreen, Julie Laplace, Søren L. Johansen, Julie K. T. Sørensen, Mohamed Chamlali, Martin W. Berchtold, Søren T. Christensen, Zeinab Anvarian, Helen L. May-Simera, Lotte B. Pedersen

**Author notes:** Corresponding author: Lotte B. Pedersen. These authors contributed equally (Csenge K. Rezi, Alina Frei, Fabiola Campestre). Corresponding author: Helen May-Simera. Further information and requests for resources and reagents should be directed the lead contact Lotte B. Pedersen.

## Abstract

Dynamic control of ciliary membrane protein content is crucial for the organelle’s homeostasis and signaling function and involves removal of ciliary components by BBSome-mediated export, endocytic retrieval and/or extracellular vesicle (EV) shedding. We report that KIF13B regulates ciliary protein composition and EV shedding in cultured kidney epithelial cells, with effects that vary over time. In early stages of ciliation *Kif13b*^-/-^ cells aberrantly accumulate PC2, FLOT1, and HGS within cilia. These cells also produce fewer small EVs through the GW4869-sensitive, nSMase2 pathway, and release large EVs enriched with CCDC198 and the centriole distal appendage protein CCDC92, which also localizes to the ciliary tip. Upon cilia maturation, *Kif13b*^-/-^ cells accelerate large EV release of numerous ciliary proteins, including PC2, BBSome components, and IFT proteins, which correlates with gradual depletion of CCDC92 and PC2 from the ciliary tip and shaft, respectively. Furthermore, over time, *Kif13b*^-/-^ cells show an upregulation in the release of small EVs, which differ in composition from wild-type small EVs. Specifically, the mutant small EVs lack several proteins that are enriched in small EVs from BBSome-deficient cells, such as the palmitoyl transferase ZDHHC5, which localizes to cilia, accumulates within cilia of BBSome-deficient cells, and regulates ciliary length and PC2 levels. Collectively, our work suggests that KIF13B acts at the level of centriole distal appendages to limit ciliary protein entrance and promote endocytic retrieval downstream of the BBSome. Furthermore, this study shows for the first time that CCDC198 and ZDHHC5 localize to primary cilia, suggesting they are potential novel ciliopathy candidates.

## Introduction

Primary cilia are antenna-like sensory organelles that protrude from the surface of many vertebrate cell types, including polarized kidney epithelial cells, and play important roles in coordinating a range of developmental and homeostatic signaling pathways that regulate cell behavior, cell cycle progression and differentiation ^1,2^. Mutations in genes that regulate cilia assembly, maintenance, composition or function are causative of pleiotropic diseases called ciliopathies, which include several diseases with renal phenotypes, e.g. nephronophthisis (NPHP), Bardet-Biedl Syndrome (BBS) and polycystic kidney disease (PKD) ^2,3^.

The ciliary membrane, which ensheathes the microtubule-based axoneme, is enriched for specific proteins (e.g. receptors and ion channels) and lipids (e.g. phosphoinositides and sphingolipids) that function in various signaling pathways ^4,5^. For example, in kidney epithelial cells the transmembrane proteins polycystin-1 (PC1) and polycystin-2 (PC2), which are encoded by the autosomal dominant (AD) PKD genes *PKD1* and *PKD2*, respectively ^6,7^, are concentrated in primary cilia ^8,9^. Here they form a calcium-permeable nonselective cation channel complex required for maintaining normal renal tubular structure and function. Mutations in *PKD1*, *PKD2* or other genes that impair ciliary localization of PC1 or PC2 are associated with ADPKD, indicating that appropriate regulation of ciliary targeting and homeostasis of the PC1-PC2 complex is critical for kidney development, homeostasis and function ^10–12^.

In polarized kidney epithelial cells, targeting of transmembrane proteins from their site of synthesis in the ER/Golgi to the primary cilium is complex, and the specific vesicular trafficking route employed can vary from protein to protein ^13,14^. For example, post-Golgi vesicles carrying PC2 are thought to be transported via recycling endosomes and/or the plasma membrane prior to reaching the base of the primary cilium where the vesicles dock at the basal body transition fibers; PC2 then crosses the transition zone and is imported into the cilium ^11,13,15^. Following import into the cilium, additional mechanisms are employed to down-regulate ciliary PC2 levels and ensure organelle homeostasis, including ciliary retrieval and endocytic sorting, as well as release of PC2-containing extracellular vesicles (EVs) ^11^.

Extracellular vesicles are released by many cell types and mediate intercellular transport of different macromolecules to regulate various physiological and pathological processes. In general, two major subtypes of EVs have been categorized based on their origin: exosomes, which are small (ca. 50-150 nm in diameter) and derived from the multivesicular body (MVB), and ectosomes (also known as microvesicles), which vary in size from ca. 100 nm to several micrometers in diameter and arise from outward protrusions of the plasma membrane that are subsequently excised and released into the environment ^16,17^. Both primary and motile cilia are known to release EVs, which may serve to regulate ciliary membrane homeostasis and intercellular signaling ^18–21^. It is thought that EVs derived from cilia are ectosomes, generated by outward budding of the ciliary membrane ^18^, which can range in size and content depending on whether they are released from the ciliary tip or base ^22^. Hereafter, we designate exosomes and small ectosomes as small EVs, and larger ectosomes as large EVs.

The cellular origin of specific EV subtypes can be difficult to distinguish in biochemical preparations, and similar EV cargo sorting and biogenesis pathways may be employed dynamically to generate diverse subpopulations of small and large EVs in a context-dependent manner ^16,17^. The specific mechanism involved in EV cargo sorting and biogenesis is thought to be dictated by the cargo, and at least four pathways have been described: (i) the ESCRT pathway involving HGS- and STAM-mediated sorting of ubiquitylated cargoes; (ii) the ALIX-syntenin pathway that bypasses the early ESCRT machinery; (iii) the GW4869-sensitive, wedge-driven curvature pathway involving neutral sphingomyelinase 2 (nSMase2)/ceramide-dependent EV formation of cargoes concentrated in lipid rafts enriched for Flotillins (FLOT1, FLOT2) and cholesterol; and (iv) actin-mediated formation of membrane blebs to generate ectosomes or large EVs ^17^. A subtype of the latter includes the ARMMs pathway (arrestin domain-containing protein 1-mediated microvesicles), which is facilitated by the ITCH ubiquitin E3 ligase, the palmitoyl transferase ZDHHC5 and the ESCRT-1 protein TSG101, amongst others ^23,24^.

While the mechanisms involved in ciliary EV cargo sorting and biogenesis are only beginning to be uncovered ^19,21^, it is well established that components of the actin cytoskeleton promote EV release from cilia ^25–29^, whereas the BBSome, a complex of BBS proteins that functions as a cargo adaptor for ubiquitinated membrane proteins during retrograde intraflagellar transport (IFT) ^30–32^, suppresses the process, both during homeostatic and signal-dependent EV release ^4,25,29,33,34^. For example, combined depletion of BBS4 and BBS5 from hTERT-immortalized human retinal pigment epithelial (RPE1) cells or *Caenorhabditis elegans* sensory neurons caused aberrant ciliary accumulation of the EV cargo PC2 due to impaired retrieval of ubiquitinated PC2 from cilia ^35^. Moreover, studies in *C. elegans* male sensory neurons indicated that downregulation of ciliary polycystins involves the ESCRT-0 proteins HGS and STAM, which function on early endosomes to direct the PC1-PC2 complex for lysosomal degradation ^36^. This pathway may also involve RAB5 and caveolin-1 (CAV1), which together with endosome maturation factors Rabenosyn-5 and VPS45 promote endocytic sorting of PC2 at the periciliary membrane compartment ^35,37^. Apart from employing BBSome-mediated retrieval and endocytic sorting to regulate ciliary polycystin homeostasis, *C. elegans* sensory neuronal cilia also exhibit environmental release of polycystin-containing EVs via a mechanism involving the kinesin-3 motor protein KLP-6; in *klp-6* mutant animals such EVs are not released into the environment causing excessive shedding of EVs into the glial lumen surrounding the ciliary base ^38^. The release of polycystin-containing EVs from cilia is evolutionarily conserved as polycystin homologs were detected in cilia-derived EVs purified from the green alga *Chlamydomonas reinhardtii* ^39,40^ and in EVs isolated from human urine ^41^ as well as the mouse embryonic node ^42^. However, the mechanisms underlying ciliary polycystin homeostasis and EV release in vertebrates remain incompletely understood.

We previously showed that a mammalian homolog of *C. elegans* KLP-6, the kinesin-3 motor protein KIF13B, localizes to primary cilia of RPE1 cells where it mediates formation of a CAV1-enriched membrane domain at the ciliary base, thereby regulating Sonic hedgehog signaling ^43^. Furthermore, we found that KIF13B undergoes transient and burst-like movement within primary cilia of RPE1 cells and is occasionally released from cilia in EV-like particles ^44^.

In this study, we demonstrate that KIF13B regulates ciliary protein composition and EV release in mouse cortical collecting duct (mCCD) cells in a time-dependent manner. In the early stages of ciliation, *Kif13b*^-/-^ cells accumulate excessive amounts of PC2, FLOT1, and HGS within cilia, produce fewer small EVs through the GW4869-sensitive, nSMase2 pathway, and aberrantly release large EVs enriched for CCDC198 (also known as FAME; Factor Associated with Metabolism and Energy), which we show localizes to cilia, as well as for the centriole distal appendage protein CCDC92, which we find also localizes to the ciliary tip. Upon prolonged serum depletion to induce cilia maturation, large EV content of numerous ciliary proteins such as PC2, BBSome and IFT components is upregulated in the *Kif13b*^-/-^ cells, and this is correlated with gradual depletion of PC2 from the ciliary compartment and loss of CCD92 from the ciliary tip. In addition, *Kif13b*^-/-^ cells also upregulate the rate of small EV release over time, but these EVs differ in composition from wild-type (WT)-derived small EVs by lacking several proteins that are conversely enriched in small EVs from BBSome-deficient cells. These include the palmitoyl transferase ZDHHC5, which we show localizes to cilia, accumulates in cilia upon BBSome inactivation, and controls ciliary length and PC2 levels.

Our findings offer new insights into the complex and dynamic processes underlying ciliary protein homeostasis and EV release, and position KIF13B as a pivotal regulator of these mechanisms. Moreover, our study demonstrates for the first time that CCDC198 and ZDHHC5 localize to primary cilia, identifying them as putative, novel ciliopathy candidates.

## Results

### KIF13B loss causes time-dependent changes in ciliary PC2 levels

In *C. elegans*, kinesin-3 motor KLP-6 regulates ciliary homeostasis and EV shedding of PKD-2 ^38,45,46^. To test if KIF13B similarly regulates ciliary PC2 homeostasis and EV shedding, we knocked it out in mCCD cells and confirmed the absence of KIF13B in *Kif13b*^-/-^ cells by western blotting (Fig. 1a). Consistent with previous work ^47,48^ mCCD cells formed primary cilia upon serum starvation for 24 or 72 h, and loss of KIF13B did not affect ciliation frequency under these conditions (Fig. S1a, b). Total cellular levels of PC2 in *Kif13b*^-/-^ cells were comparable to those of WT cells, both for cells grown in the presence of serum (+FBS) or depleted for serum for 24, 48, or 72 h (Fig. 1 a, b; Fig. S1c). We then analyzed the ciliary localization of PC2 in WT and *Kif13b*^-/-^ cells by immunofluorescence microscopy (IFM). Using a previously described anti-PC2 antibody ^49^ to stain for PC2 in 24 h serum-starved cells, we observed significant accumulation of PC2 within cilia of *Kif13b*^-/-^ cells, as compared to WT cells (Fig. 1c, d). Stable expression of mCherry-KIF13B in the *Kif13b*^-/-^ cells (Fig. S1d, e) rescued this phenotype (Fig. 1c, d), and similar results were obtained using an alternative antibody against PC2 (Fig. 1e, f). In contrast, IFM analysis of 72 h serum-starved cells showed that ciliary PC2 levels in *Kif13b*^-/-^ mutant cells were significantly lower than for WT cells, a phenotype that was rescued by stable expression of mCherry-KIF13B (Fig. 1e, f). Thus, KIF13B regulates ciliary PC2 levels in a time-dependent manner.

**Figure 1.**
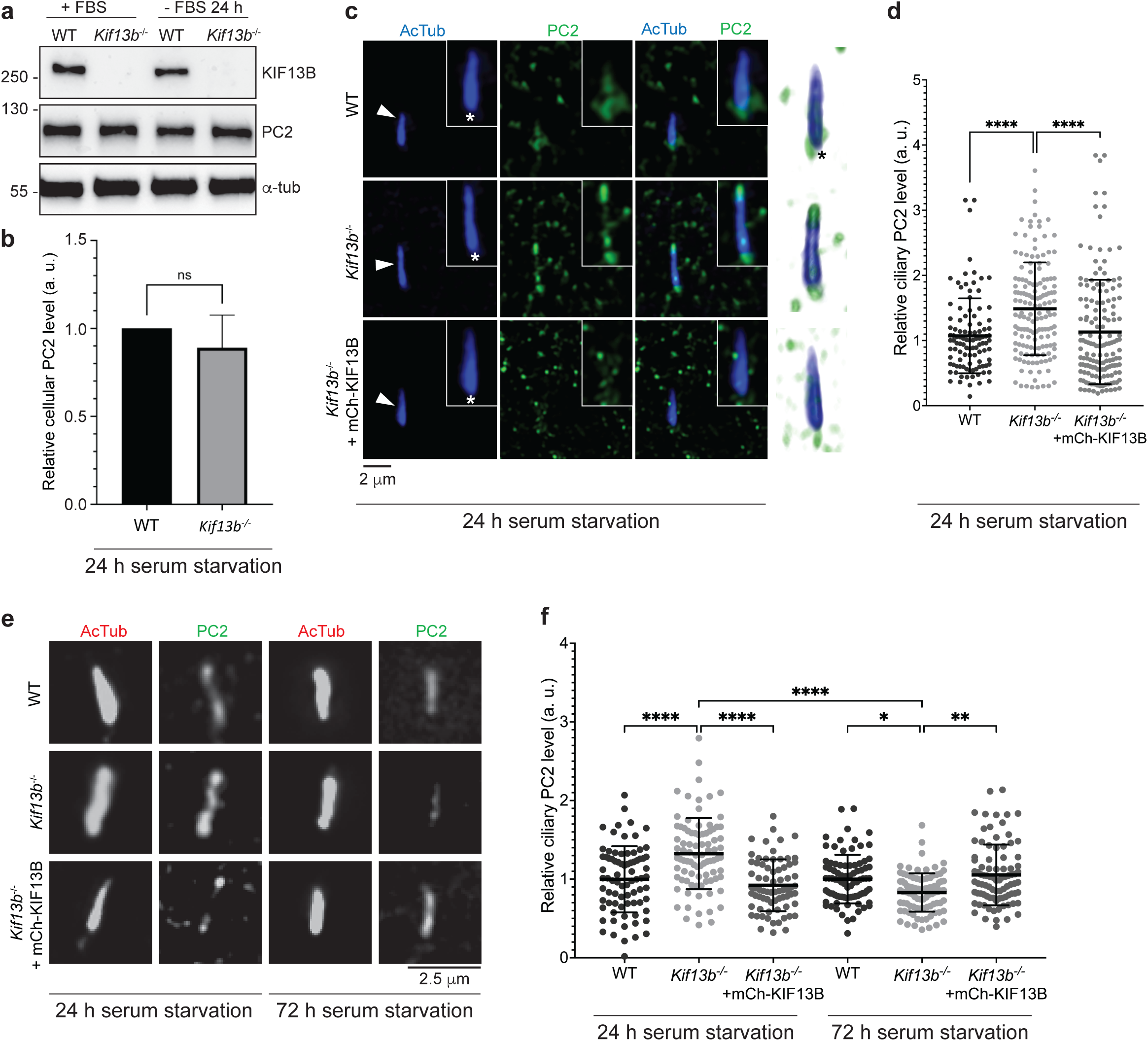
Ciliary PC2 localization in WT, *Kif13b*^-/-^ and rescue mCCD cell lines. (**a**) Western blots of WT and *Kif13b*^-/-^ mCCD cells grown with (+) or without (-) FBS for 24 h; blots were probed with antibodies as indicated; α-tubulin (α-tub) serves as loading control. Molecular mass markers are shown in kDa to the left. (**b**) Quantification of relative cellular PC2 levels in WT and *Kif13b*^-/-^ cells after 24 h of serum starvation, based on western blots as shown in panel a (n=3). (**c**) IFM analysis of indicated mCCD cell lines after 24 h of serum starvation. Acetylated α-tubulin (AcTub; blue) marks the ciliary axoneme (arrowheads) and PC2, stained with an antibody described in ^49^, is shown in green. The ciliary base is marked with asterisk. Insets show enlargement of the cilium-centrosome axis, and the rightmost panels are 3D renderings thereof. mCh-KIF13B: mCherry-KIF13B. (**d**) Quantification of relative ciliary PC2 staining intensities of the indicated mCCD cell lines, based on images as shown in (c). (**e**) IFM analysis of indicated mCCD cell lines following serum starvation for 24 or 72 h and using commercially available antibodies against PC2 and acetylated α-tubulin (AcTub). (**f**) Quantification of relative ciliary PC2 staining intensity in the indicated mCCD cell lines, based on images as shown in (e). For quantifications in (d, f) background-corrected mean fluorescence intensity (MFI) of PC2 was measured for 50 cilia per cell line per experiment and normalized to that of WT 24 h mean value (n=3). *, p<0.05; **, p<0.01; ****, p<0.0001; ns, not significant.

### KIF13B loss causes increased membrane bulging from the side of cilia

The elevated ciliary PC2 levels observed in the 24 h serum-starved *Kif13b*^-/-^ cells could result from increased ciliary entrance of PC2, or impaired removal of PC2 from cilia via EV release or by retrograde transport and endocytic retrieval. Release of EVs from cilia is associated with ciliary membrane bulging ^25^ whereas retrieval and endocytosis have been linked to the ciliary pocket, an invagination of the periciliary membrane ^50–53^. To assess if loss of KIF13B affects ciliary membrane bulging and pocket formation, we analyzed ciliary morphology in 24 h or 72 h serum-deprived WT, *Kif13b*^-/-^ and rescue lines by scanning electron microscopy (SEM). We found that average ciliary lengths were about 2 μm for all cell lines, with a tendency for longer cilia in the 72 h serum-starved *Kif13b*^-/-^ cells (Fig. 2a, b; Fig. S2a). Approximately 50-60% of the cilia displayed a pocket, regardless of cell line and starvation conditions (Fig. 2a, b; Fig. S2b), and ciliary membrane bulging was similarly observed in all conditions (Fig. 2a-d). However, whereas cilia with membrane bulges at the tip were detected at similar frequency in the 72 h serum-starved WT and *Kif13b*^-/-^ cells, membrane bulging at the side of cilia was significantly more frequent in *Kif13b*^-/-^ cells (Fig. 2d). A similar trend was observed for the 24 h serum-starved cells, although the difference was not statistically different (Fig. 2c). The size of the membrane bulges observed at the tip and side of cilia in the different mCCD cell lines (Fig. 2a, b) is equivalent to or larger than the diameter of the cilium (100 nm), classifying them in the size range of large EVs ^16^. These membrane bulges resemble budding EVs previously observed in primary cilia of murine inner medullary collecting duct (IMCD)3, primary biliary epithelial and neuroepithelial cells, as well as Madin-Darby canine kidney (MDCK) and hTERT-RPE1 cells ^25,41,54–58^. Thus, loss of KIF13B has no major influence on ciliary length or pocket formation in mCCD cells but leads to increased membrane bulging from the side of cilia.

**Figure 2.**
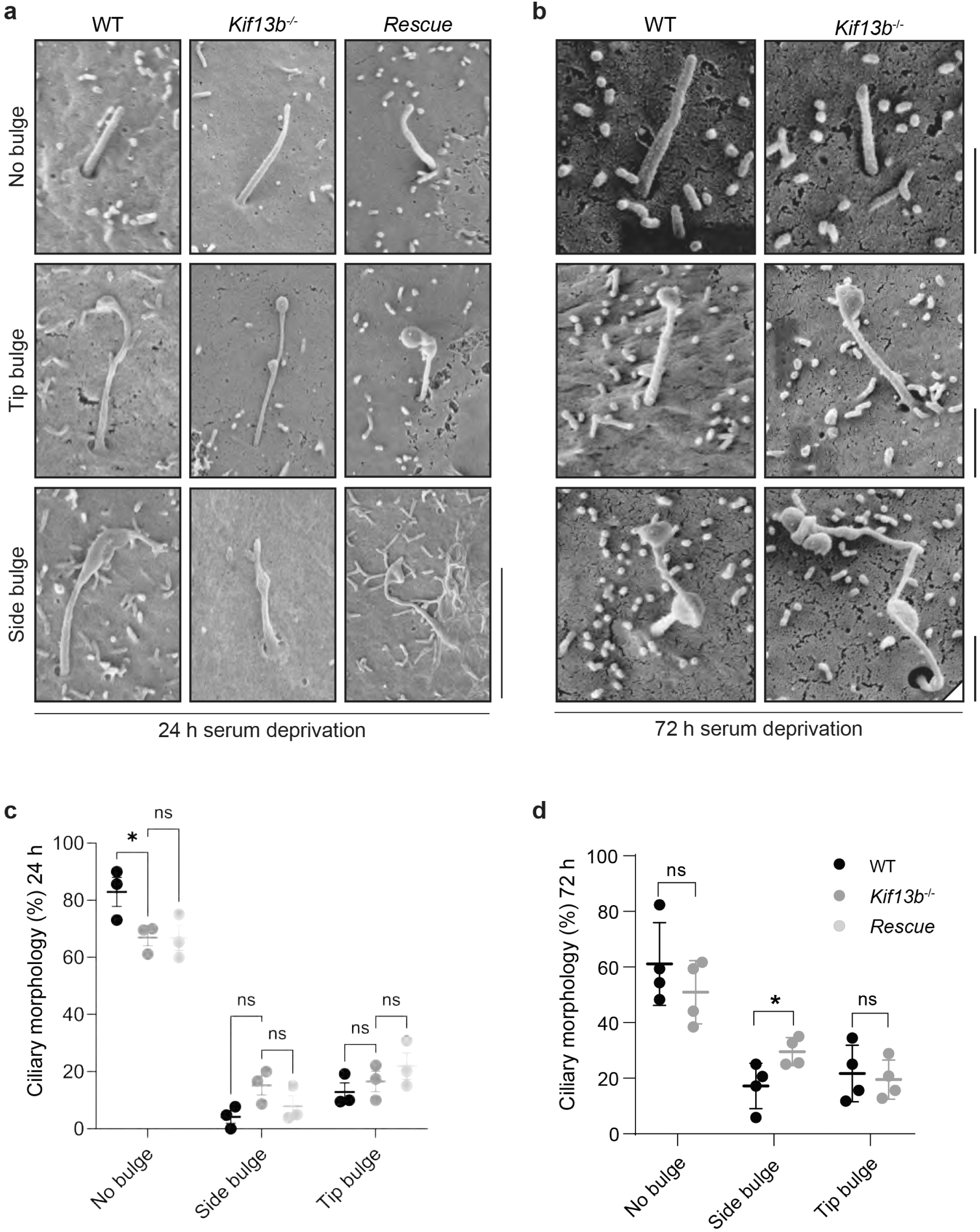
Ciliary morphology of mCCD cell lines. (**a, b**) Representative images of different cilia morphologies observed by SEM in the indicated mCCD cell lines following 24 h (**a**) or 72 h (**b**) of serum starvation. Scale bars: 3 μm in panel a, 1 μm in panel b. (**c**) Quantification of ciliary membrane bulges in 24 h serum starved cells, with 61-67 cilia analyzed in total per cell type (n=3). (**d**) Quantification of ciliary membrane bulges in 72 h serum starved cells, with 29-77 cilia analyzed per cell line per experiment (n=4; 163 and 208 cilia analyzed in total for WT and *Kif13b*^-/-^ cells, respectively). *, p<0.05; ns, non-significant.

### Dynamic ciliary localization and EV release of KIF13B in mCCD cells

We investigated if KIF13B localizes within cilia of mCCD cells by performing live cell imaging of cells stably expressing mNeonGreen (mNG)-KIF13B in the *Kif13b*^-/-^ background (Fig. S1d, e) and subjecting the cells to 24-72 h of serum starvation followed by SiR-tubulin staining to visualize the ciliary axoneme. In cells serum-starved for 24-48 h mNG-KIF13B transiently accumulated and moved bidirectionally within 21% of cilia analyzed (n=72; Movie 1), reminiscent of our previous observations in ciliated hTERT-RPE1 cells transiently expressing eGFP-KIF13B ^44^. Similar observations were made for cells serum-starved for 72 h, where mNG-KIF13B transiently accumulated and moved within 32% of cilia analyzed (n=57). In addition, under these conditions mNG-KIF13B occasionally seemed to be released from the ciliary tip (Movie 2), as seen previously in hTERT-RPE1 cells ^44^. Consistent with the latter observation, western blot analysis of large and small EVs purified from spent medium of 72 h serum-starved mCCD cell cultures by centrifugation (large EVs) and lectin-mediated precipitation (small EVs) ^59^, respectively (Fig. S3a), identified KIF13B predominantly in large EVs, whereas ciliary proteins like ARL13B, acetylated α-tubulin and PC2 were detected in both large and small EV samples under these conditions (Fig. S3b, c). Notably, loss of KIF13B did not prevent EV release of PC2 but seemed to shift its relative distribution between large and small EV populations (Fig. S3b, c; see also below). We conclude that KIF13B localizes to cilia of mCCD cells and that both KIF13B and PC2 are released in EVs from these cells.

### Multiple ciliary proteins accumulate in large EVs from *Kif13b*^-/-^ cells over time

To further explore how KIF13B affects EV release of PC2 and/or other ciliary proteins, we next subjected large EVs, purified from 24 or 72 h serum-starved WT and *Kif13b*^-/-^ cells by centrifugation (Fig. 3a), to mass spectrometry (MS) analysis to assess their protein content. Proteins that were significantly altered in abundance between WT and *Kif13b*^-/-^ samples were divided into two Tiers based on stringency criteria (see Materials and Methods for details). For the 24 h samples, we identified seven Tier1 proteins that were significantly altered in abundance in large EVs from the *Kif13b*^-/-^ cells as compared to WT large EVs, four of which (CTPS2, POFUT2, DCTN6, NHLC2) were depleted from the mutant-derived EVs, the remaining three (CCDC198, CCDC92, MLF2) being enriched in mutant-derived EV samples compared to controls (Fig. 3b; Table S1). For the 72 h large EV samples we identified 72 Tier 1 proteins that were significantly altered in abundance between *Kif13b*^-/-^ and WT samples, including numerous ciliary proteins such as IFT and BBSome components (Fig. 3c; Table S2). Consistently, Gene Ontology (GO) analysis of the Tier 1 and Tier 2 proteins in the latter, focusing on the cellular component category, identified “intraciliary transport particle B binding” as the most highly enriched GO term in this dataset, followed by the GO terms “tubulin binding” and “cytoskeletal binding” (Fig. 3d; Table S5). Similarly, GO analysis for the molecular function category identified “intraciliary transport particle” and “cilium” as the most highly enriched GO terms among this dataset (Fig. 3e; Table S5). Of note, both PC2 and PC1 were also significantly enriched in the 72 h large EV samples from *Kif13b*^-/-^ cells compared to WT (Tier 2 hits; Table S2) but were either not detected (PC1) or unchanged in relative abundance (PC2) in the 24 h large EV samples from WT and *Kif13b*^-/-^ cells. Thus, despite the presence of KIF13B in large EVs (Figure S3b, c), these results suggests that KIF13B does not promote large EV release of PC2 from cilia, but rather suppresses this process. Since ciliation frequency (Figure S1b) and ciliary length (Figure S2a) were similar between WT and *Kif13b*^-/-^ cells, we conclude that loss of KIF13B causes increased release of ciliary proteins in large EVs over time without compromising overall structural ciliary integrity.

**Figure 3.**
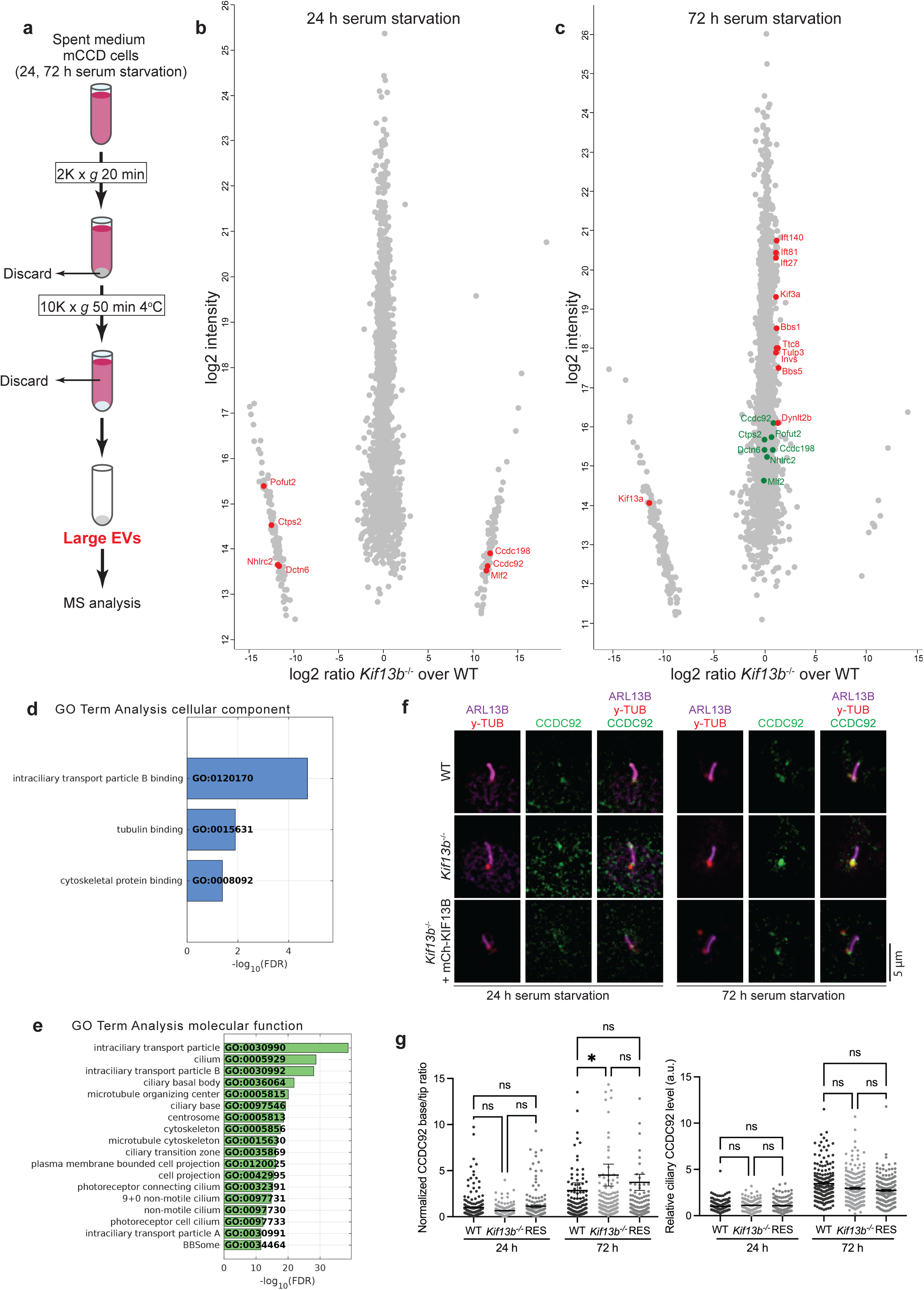
Altered composition of large EVs from *Kif13b*^-/-^ cells. (**a**) Purification scheme of large EVs used for MS analysis. (**b**, **c**) Scatterplots comparing the protein content of large EVs isolated from WT and *Kif13b*^-/-^ mCCD cells after 24 h (**b**) and 72 h (**c**) of serum starvation, based on MS analysis of 5 experimental replicates per condition. Selected proteins that differ significantly in abundance between mutant and WT samples are highlighted in red. (**d**, **e**) GO Term Analysis for the categories cellular component (**d**) and molecular function (**e**) for proteins significantly altered in abundance in large EVs of *Kif13b*^-/-^ mCCD cell cultures after 72 h of serum starvation (Table S2, Table S5). The tables show the GO terms that are significantly enriched (Fisher’s exact test value ≤0.05), listed according to their enrichment ratio. (**f**) IFM analysis of CCDC92 in the indicated cell lines. Cells were co-stained for ARL13B and γ-tubulin (γ-tub) to label the ciliary membrane and centrioles, respectively. (**g**) Quantification of ciliary base to tip ratio (left) or total ciliary staining intensity (right) of CCDC92 in the indicated cell lines, based on images as shown in (f). Background-corrected MFI of CCDC92 was measured for 40-60 cilia per cell line per experiment and normalized to that of WT 24 h mean value (n=3). *, p<0.05; ns, not significant.

### CCDC92 and CCDC198 localize to the ciliary base and tip

To investigate the mechanism by which KIF13B may affect large EV release from cilia, we initially focused our attention on CCDC92 and CCDC198, both of which were significantly enriched in large EVs from 24 h serum-starved *Kif13b*^-/-^ cells compared to WT samples (Fig. 3b; Table S1). CCDC92 is a centrosomal distal appendage protein (DAP) that interacts with the DAPs CEP164 and TTBK2 ^60–63^, which in turn binds to CSNK2A1 that plays a crucial role in controlling ciliary protein trafficking and suppresses ciliary EV release in mouse embryonic fibroblasts ^63^. CCDC198 (also known as FAME) is a relatively poorly studied protein linked to mammalian energy balance and kidney physiology ^64^, which interacts directly with FNBP1L, a CDC42 effector that promotes actin-based endocytosis downstream of CDC42 ^64,65^. We confirmed by IFM analysis that CCDC92 localizes to the ciliary base of mCCD cells, where it additionally concentrates at the ciliary tip (Fig. 3f). Interestingly, while total ciliary levels of CCDC92 were unchanged in the *Kif13b*^-/-^ cells relative to controls, both after 24 and 72 h of serum starvation, the ciliary base to tip ratio of CCDC92 initially seemed to be decreased in the mutant cells, a phenotype that was completely reversed at the 72 h time point (Fig. 3f, g). Combined with our large EV MS data (Fig. 3b, Table S1) these results indicate that KIF13B promotes retention of CCDC92 at the ciliary base, and that loss of KIF13B causes CCDC92 to accumulate aberrantly at the ciliary tip from where it is released in large EVs and eventually depleted. In support of this, by IFM analysis of our *Kif13b*^-/-^ cells we occasionally observed CCDC92 in EV-like particles that were positive for the ciliary marker acetylated tubulin (Fig. S4c). We also performed similar IFM analysis of CCDC198 and found that it localizes to the plasma membrane, as expected ^64^, but also to the primary cilia where it was mostly concentrated at the base and the tip (Fig. S4a). However, we did not detect significant changes in the overall ciliary levels of CCDC198 between *Kif13b*^-/-^ and WT cells under these conditions (Fig. S4b); for technical reasons we were unable to assess CCDC198 ciliary base to tip ratios as done for CCDC92. We conclude that KIF13B regulates the ciliary base to tip ratio of CCDC192 in a time-dependent manner, and CCDC198 localizes to primary cilia.

### KIF13B promotes GW4869-sensitive/ceramide-dependent small EV release

As our initial analysis identified ciliary proteins in both large and small EV fractions (Fig. S3b, c), we next investigated the effect of KIF13B loss on small EV release. We collected spent medium from WT and *Kif13b*^-/-^ mCCD cell cultures following 24, 48 or 72 h of serum starvation and used differential centrifugation ^20^ to purify small EVs (Fig. 4a). Successful purification of small EVs was confirmed by transmission electron microscopy (TEM), which revealed the characteristic presence of spherical biconcave structures 100-200 nm in size (Fig. 4b), and by western blot analysis using antibodies specific for common small EV marker proteins (Fig. S3d). Nanoparticle tracking analysis (NTA) showed a significant decrease in the number of small EVs released from 24 h serum-deprived *Kif13b*^-/-^ cultures compared to WT controls (Fig. 4c). Intriguingly, prolonged serum-deprived conditions (72 h in total) caused a significant increase in the relative rate of small EVs release in the *Kif13b*^-/-^ cells bringing the rate of release comparable to WT levels, which compensated the difference at baseline release after 24 h (Fig. 4c, d). This suggests an adaptive mechanism in small EV release in the *Kif13b^-/-^*cells over time. Treatment of the 24 h serum-starved cultures with GW4869, a well-established inhibitor of nSMase2/ceramide-dependent EV biogenesis ^17^, led to an overall reduction in small EV numbers for both cell lines, as expected, suggesting that a significant proportion of small EVs are generated by an nSMase2/ceramide-dependent pathway in these cells (Fig. 4e, f). However, the inhibitory effect of GW4869 treatment on the number of small EVs released from WT cultures was significantly higher compared to that of *Kif13b*^-/-^ cultures (Fig. 4e, f), indicating that the *Kif13b*^-/-^ cells are already impaired in small EV biogenesis or release via this pathway.

**Figure 4.**
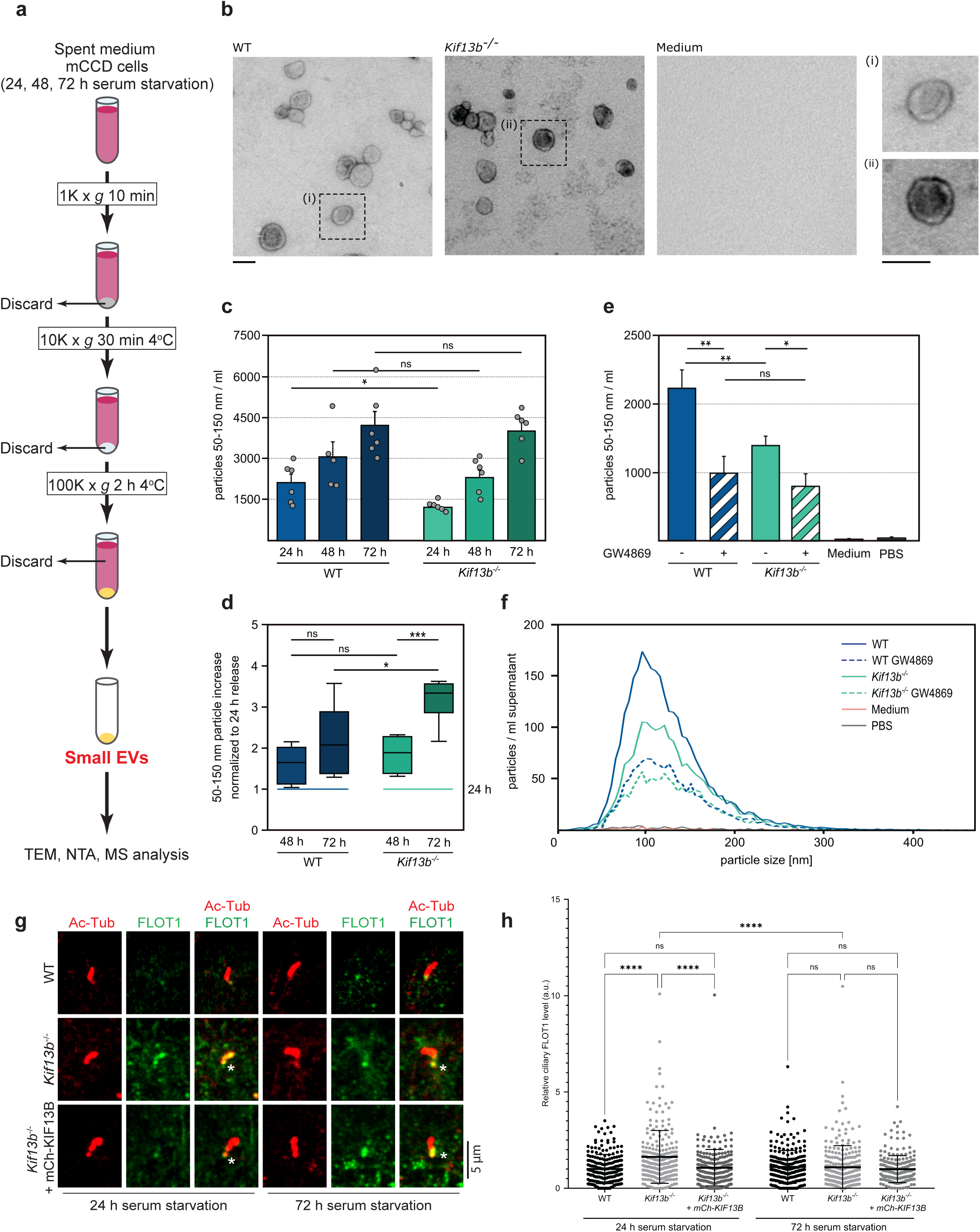
*Kif13b*^-/-^ mCCD cells are defective in nSMase2/ceramide-dependent small EV release. (**a**) Scheme of procedure used for purification of small EVs from spent cell culture medium. (**b**) Negative staining and TEM analysis of small EVs purified from spent cell culture medium after 48 h of serum starvation (scale bars, 100 nm). (**c**) NTA profiling of small EVs from spent cell culture medium collected after 24, 48, and 72 h of serum starvation, respectively. Data are represented as mean values; error bars show SEM (n=5-6 per cell line per condition). (**d**) Relative increase in small EV release of WT and *Kif13b*^-/-^ cultures at time point 48 and 72 h serum starvation, respectively, compared to the 24 h time point. Based on data shown in (c). (**e**, **f**) NTA profiling of small EVs purified from spent medium of 24 h serum-deprived WT and *Kif13b*^-/-^ mCCD cell cultures incubated with (+) or without (-) 10 μM GW4869 for 24 h prior to collection of the spent medium (n=14 for untreated; n=7 for GW4869; n=3 for Medium; n=10 for PBS). Data are represented as mean values of particles 50-150 nm in size. Error bars in (e) show SEM. For (c-e), statistics was done using Shapiro-Wilk test and Unpaired t-test, two-tailed. *, p<0.05; **, p<0.01; ***, p<0.001; ****, p<0.0001; ns, non-significant. (**g**) IFM analysis of indicated mCCD cell lines after 24 or 72 h of serum starvation, stained for acetylated α-tubulin (AcTub; red) and FLOT1 (green). The ciliary base is marked with asterisk. mCh-KIF13B: mCherry-KIF13B. (**h**) Quantification of relative ciliary FLOT1 staining intensities of the indicated mCCD cell lines, based on images as shown in (g). Background-corrected MFI of FLOT1 was measured for 40-60 cilia per cell line per experiment and normalized to that of WT 24 h mean value (n=3). ****, p<0.0001; ns, non-significant.

N-SMase2/ceramide-dependent EV biogenesis involves localization of cargoes to lipid rafts enriched in Flotillins (FLOT1, FLOT2) and cholesterol ^17^. The ciliary membrane is highly enriched in ceramides ^66,67^, KIF13B binds directly to the hydrophobic ceramide backbone of glycosphingolipids ^68^ and is required for establishing a membrane microdomain at the ciliary base of hTERT-RPE1 cells, enriched for the cholesterol-binding CAV1 protein and FLOT2 ^43^. Moreover, CAV1, FLOT1 and FLOT2 were identified as proximity interactors of the basal body DAP TTBK2 in HEK293T cells ^63^, and FLOT1 and FLOT2 were reported to localize to the ciliary base of odontoblasts and kidney epithelial cells where they interacted with OFD1, PC1 and PC2 ^69^. We therefore tested if loss of KIF13B affects ciliary FLOT1/2 localization. In support of this, IFM analysis of 24 h serum-starved mCCD cells using FLOT1-specific antibody showed that *Kif13b*^-/-^ cells aberrantly accumulate FLOT1 within cilia, and stable expression of mCherry-KIF13B rescued this phenotype (Fig. 4g, h); the total cellular level of FLOT1 was not affected by loss of KIF13B (Fig. S3e). Notably, in 72 h serum-starved *Kif13b*^-/-^ cells the ciliary/basal body level of FLOT1 was the same as for WT cells (Fig. 4g, h), indicating that KIF13B regulates ciliary levels of FLOT1 in a time-dependent manner similar to PC2.

### KIF13B regulates small EV protein content

To determine if loss of KIF13B affects the protein content of small EVs, we also analyzed small EV samples from 24 and 72 h serum-starved WT and *Kif13b*^-/-^ cultures by MS. For the 24 h small EV samples three proteins, THBS1, Vimentin and HGS, were significantly depleted from the *Kif13b*^-/-^ samples compared to WT (Fig. 5a; Table S3); the depletion of THBS1 from the mutant small EVs could be due to altered vesicular trafficking of POFUT2 in the *Kif13b*^-/-^ cells (Fig. 3b; Table S1), since POFUT2 catalyzes the addition of O-fucose to THBS1 ^70^. For the 72 h samples 52 proteins were significantly altered in the *Kif13b*^-/-^ sample compared to WT, with the majority being depleted from the mutant-derived EVs (Fig. 5b, d; Table S4). GO analysis of the latter proteins, focusing on the cellular component category, identified “cytoplasm” and “intracellular anatomical structure” as the most highly enriched GO terms in this dataset (Fig. 5c; Table S5). GO analysis focusing on molecular function ‘only’ identified protein binding (Table S5). Specific proteins depleted from the 72 h mutant EV samples included TTC8, a BBS protein ^71^ and known BBSome component ^30^; the acid ceramidase ASAH2; the deubiquitinase STAMBP, which regulates cell surface receptor trafficking at ESCRT-0 positive endosomes by directly binding to the HGS-STAM complex and regulating its ubiquitination status ^72^; the ubiquitin E3 ligase ITCH that binds and ubiquitylates several proteins including HGS and STAM ^73^; and the palmitoyltransferase ZDHHC5 (Fig. 5b; Table S4). PC1 and PC2 were also detected in our small EV proteomes of 72 h serum-starved WT and *Kif13b*^-/-^ cultures but were unchanged in relative abundance in the mutant small EVs compared to WT under these conditions. We did not detect PC1 nor PC2 in small EVs from 24 h serum-starved cultures (Table S3). Interestingly, we noted that several of the differentially regulated small EV proteins detected in our MS analysis, including THBS1, HGS and ZDHHC5, were identified in the ciliary proteome of IMCD3 cells, and HGS accumulated in cilia upon inhibition of actin polymerization with cytochalasin D ^74^. Moreover, HGS, as well as 11 of the proteins that were depleted from the 72 h *Kif13b*^-/-^ small EV samples were enriched in small EVs purified from *Bbs4* and/or *Bbs6* mutant kidney medullary (KM) cell lines (Fig. 5d) ^20^. Based on these observations, we surmised that KIF13B may function downstream of the BBSome to promote endocytic retrieval of HGS and other small EV cargoes from cilia, ultimately leading to their small EV release via MVBs. To test this, we initially focused on HGS, given its known role in regulating ciliary PC2 homeostasis ^36^ and because HGS interacted directly with FLOT1 in an *in vitro* pull-down assay ^75^. By IFM analysis of WT, *Kif13b*^-/-^ and rescue mCCD lines using an antibody specific for HGS we confirmed that HGS localizes to cilia of these cells, primarily at the ciliary base (Fig. 5e). Moreover, HGS accumulated significantly within cilia of the *Kif13b*^-/-^ cells compared to the WT and rescue line, both after 24 and 72 h of starvation (Fig. 5f), but total cellular levels of HGS did not appear to be altered in the mutant cells (Fig. S3e). To assess how ciliary HGS localization is affected by loss of BBSome function, we analyzed WT and *Ift27*^-/-^ IMCD3 cells ^76^ by similar approaches, and observed significant ciliary enrichment of HGS in the *Ift27*^-/-^ IMCD3 cells compared to WT cells (Fig. 6a), whereas the total cellular level of HGS appeared to be unaffected (Fig. 6b). Interestingly, similar analysis for FLOT1 showed that it accumulates significantly at the ciliary base of *Ift27*^-/-^ IMCD3 cells as compared to WT control cells (Fig. 6c), but total cellular levels of FLOT1 were unaltered in the mutant cells (Fig. 6d). Since IFT27 connects the BBSome to IFT trains during retrograde transport out of cilia ^76,77^, these results suggest that the BBSome may facilitate ciliary export of HGS via retrograde IFT while FLOT1 is regulated by a different mechanism. Notably, IFT27 and BBS4 were found to interact with FLOT1/2 in a chemical cross-linking and yeast-two hybrid screen, respectively ^78,79^, supporting that IFT27 and the BBSome interact with Flotillins. In summary, the steady state ciliary/basal body levels of HGS and FLOT1 are negatively regulated by KIF13B and IFT27, and KIF13B promotes small EV release of HGS and several other proteins in a time-dependent manner.

**Figure 5.**
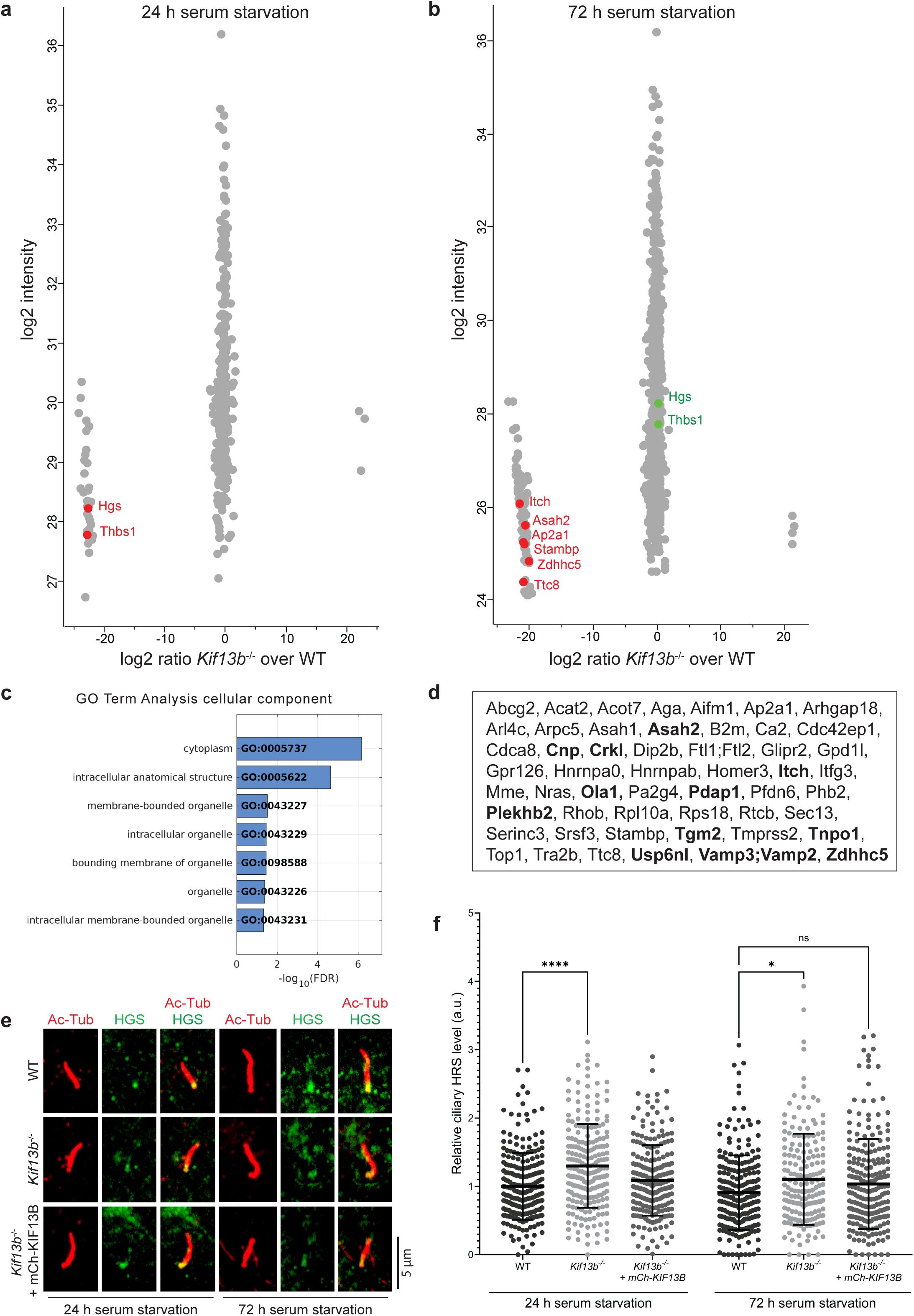
Altered composition of small EVs from *Kif13b*^-/-^ cells. (**a**, **b**) Scatterplots comparing the protein content of small EVs isolated from WT and *Kif13b*^-/-^ mCCD cells after 24 h (**a**) and 72 h (**b**) of serum starvation, based on MS analysis of 4 experimental replicates per condition. Selected proteins that differ significantly in abundance between mutant and WT samples (Tier 1 or Tier 2 hits) are highlighted in red. (**c**) GO Term Analysis for the category cellular component for proteins significantly altered in abundance in small EVs of *Kif13b*^-/-^ mCCD cell cultures after 72 h of serum starvation (Table S4, Table S5). The tables show the GO terms that are significantly enriched (Fisher’s exact test value ≤0.05), listed according to their enrichment ratio. (**d**) List of proteins significantly altered in abundance in small EVs of *Kif13b*^-/-^ mCCD cell cultures after 72 h of serum starvation (Tier 1 and Tier 2 hits). Proteins in bold were previously shown to be significantly enriched in small EVs derived from *Bbs4* and/or *Bbs6* mutant KM cells ^20^; except for TGM2, all of these proteins were depleted from the *Kif13b* mutant EVs compared to WT. (**e**) IFM analysis of indicated mCCD cell lines after 24 or 72 h of serum starvation, stained for acetylated α-tubulin (AcTub; red) and HGS (green). The ciliary base is marked with asterisk. mCh-KIF13B: mCherry-KIF13B. (**f**) Quantification of relative ciliary HGS staining intensities of the indicated mCCD cell lines, based on images as shown in (e). Background-corrected MFI of HGS was measured for 40-60 cilia per cell line per experiment and normalized to that of WT 24 h mean value (n=3). *, p<0.05; ****, p<0.0001; ns, non-significant.

**Figure 6.**
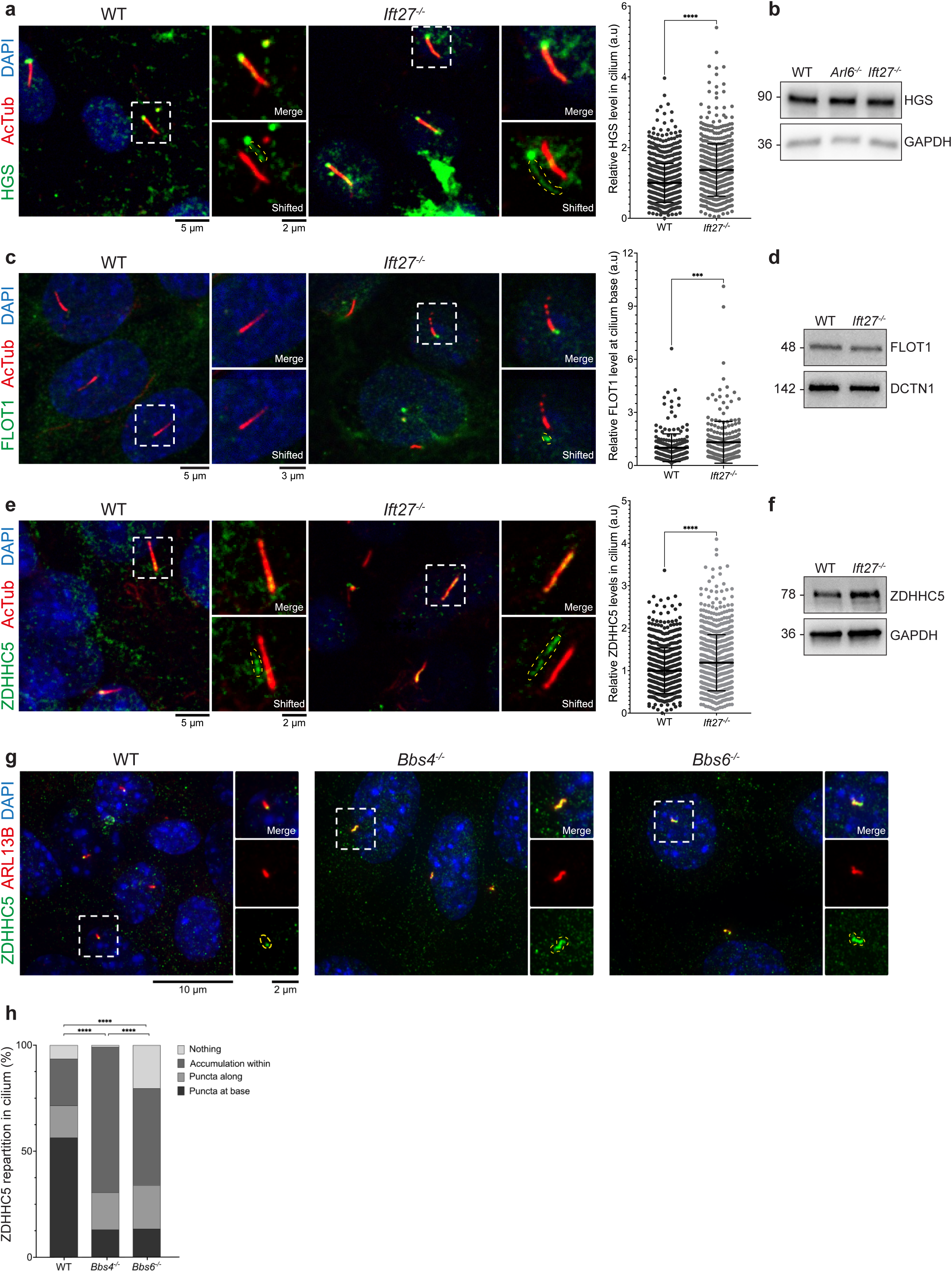
Altered ciliary localization of HGS, FLOT1 and ZDHHC5 in *Bbs* mutant cells. (**a**, **c**, **e**) Representative IFM images and quantitative analysis of ciliary staining intensities of 24 h serum-starved WT and *Ift27*^-/-^ IMCD3 cells stained with antibodies against acetylated tubulin (AcTub) and HGS (**a**), FLOT1 (**c**) or ZDHHC5 (**e**). Nuclei were stained with DAPI. Insets show merged or shifted magnifications of selected cilia (boxed regions). For quantitative analysis (right panels), the MFI of anti-HGS, -FLOT1 or -ZDHHC5 signal within the cilium (a, e) or at the ciliary base (c) was measured manually or by a semi-automated approach (see Materials and Methods) and normalized to the mean MFI for WT. For FLOT1 (c), between 40-50 cilia were measured manually per condition per experiment (n=3). ***, p<0.001; ****, p<0.0001. (**b**, **d**, **f**) Western blot analysis of EV markers in IMCD3 cells. WT and *Ift27*^-/-^ IMCD3 cells were serum-starved for 24 h and analysed by SDS-PAGE and western blotting using antibodies as indicated. GAPDH or DCTN1 were used as loading controls, and *Arl6*^-/-^ IMCD3 cell lysate included in panel b for comparison. Molecular mass markers are indicated in kDa to the left of the blots. (**g**, **h**) Representative IFM images and quantitative analysis of ciliary staining intensities of 24 h serum-starved WT, *Bbs4*^-/-^ and *Bbs6*^-/-^ KM cells stained with antibodies against ARL13B and ZDHHC5. Nuclei were stained with DAPI. Insets show merged magnifications of selected cilia (boxed regions). In panel (h), a Chi square test was used to test for contingency between sample distribution. ****, p<0.0001

### Ciliary localization and function of ZDHHC5

The above-described results indicate that KIF13B promotes small EV release by a pathway involving ceramides and Flotillins. We noted that FLOT2, and likely also FLOT1, were identified as the main palmitoylation substrates of ZDHHC5 in mouse neuronal stem cells ^80^. Since ZDHHC5 was depleted and enriched, respectively, in small EVs from 72 h serum-starved *Kif13b*^-/-^ mCCD cells (Fig. 5b; Table S4) and BBSome-deficient KM cells ^20^, we tested if ZDHHC5 is localized to primary cilia in these cells. Live imaging of WT mCCD cells stably expressing mNG-ZDHHC5 indeed showed that the latter is highly concentrated at the base and along primary cilia of these cells (Movie 3), in agreement with a previous study identifying ZDHHC5 in the ciliary proteome of IMCD3 cells ^74^. In addition, IFM staining for endogenous ZDHHC5 showed prominent localization of ZDHHC5 to the basal body and primary cilium of both WT and *Kif13b*^-/-^ mCCD cells and indicated that its ciliary levels are higher at 72 h compared to 24 h of serum starvation (Fig. S5a, b). There was a trend towards reduced ciliary levels of ZDHHC5 in the 72 h serum-starved *Kif13b*^-/-^mutant cells compared to the WT and rescue line, but this change was not statistically significant (Fig. S5b). Similar results were obtained for WT and *Kif13b*^-/-^ mutant cells stably expressing mNG-ZDHHC5 (Fig. S5c). In contrast, IFM analysis of serum-starved WT and *Ift27*^-/-^ IMCD3 cells showed that endogenous ZDHHC5 accumulates significantly within cilia of the mutant cells (Fig. 6e) without affecting overall cellular ZDHHC5 protein levels (Fig. 6f), a phenotype that was recapitulated for mNG-ZDHHC5 stably expressed in these cells (Fig. S5d) and for endogenous ZDHHC5 in *Bbs4* or *Bbs6* mutant KM cells (Fig. 6g, h).

Finally, to analyze the ciliary function of ZDHHC5 we knocked it out in mCCD cells (Fig. S6a) and analyzed WT and mutant cells by IFM with antibodies against relevant ciliary proteins. Loss of ZDHHC5 had little effect on ciliation frequency in these cells but caused a significant increase in ciliary length compared to control cells (Fig. S6b, d, e). Furthermore, ciliary levels of PC2 were significantly decreased in the *Zdhhc5*^-/-^ cells compared to WT, both after 24 and 72 h of starvation (Fig. S6b, f), but total cellular levels of PC2 were not altered in the *Zdhhc5*^-/-^ cells (Fig. S6c). In summary, ciliary ZDHHC5 levels are negatively regulated by IFT27 and the BBSome, and loss of ZDHHC5 causes ciliary lengthening and reduced ciliary content of PC2.

## Discussion

Our study shows that KIF13B regulates ciliary protein content and EV release in mCCD cells in a time-dependent manner, as summarized in Fig. 7. Specifically, *Kif13b*^-/-^ cells with newly formed cilia display significantly enriched ciliary levels of PC2, FLOT1 and HGS compared to WT cells (Fig. 7a, b), but upon prolonged serum starvation to induce cilia maturation, this phenotype is gradually reversed leading to depletion of these proteins from mutant cilia over time (Fig. 7c, d). The time-dependent changes in ciliary protein content of the *Kif13b*^-/-^ cells are paralleled by changes in the overall frequency of release and/or protein content of small and large EVs. In particular, the KIF13B-deficient cells initially release fewer small EVs than WT cells, owing to reduced nSMase2/ceramide-dependent small EV biogenesis, and the mutant EVs are depleted for HGS, THBS1 and Vimentin. Conversely, large EVs derived from 24 h serum-starved *Kif13b*^-/-^ cells are enriched for CCDC198, CCDC92 and MLF2 but depleted for several other proteins such as POFUT2 and DCTN6. Since PC2 was either not detected (small EVs) or present in equal amounts (large EVs) in EV samples from 24 h serum-starved *Kif13b*^-/-^ and WT cultures, we propose that the main reason why PC2 initially accumulates within cilia of *Kif13b*^-/-^ cells at this time point is that the mutant cells either fail to promote HGS- and Flotillin-dependent endocytic retrieval of PC2 from cilia, consistent with the known role of HGS in ciliary PC2 homeostasis in *C. elegans* ^36^, and/or have a defective/leaky diffusion barrier at the ciliary base leading to unrestricted ciliary entrance of PC2, HGS and FLOT1 (Fig. 7b). In agreement with the latter, our results suggested that KIF13B promotes retention of CCDC92 at the ciliary base, with loss of KIF13B leading to aberrant CCDC92 accumulation at the ciliary tip followed by its release in large EVs and, eventually, its depletion from the ciliary tip. Similarly, enrichment of multiple ciliary proteins such as PC2, BBSome and IFT components in large EVs of *Kif13b*^-/-^ cells after 72 h of serum starvation is likely an indirect consequence of their initial accumulation within cilia and could explain their depletion from this compartment over time (Fig. 7b, d). In agreement with this scenario, multiple studies have shown that aberrant accumulation or mis-localization of ciliary (membrane-associated) proteins in specific sub-cellular or ciliary compartments can trigger their release in EVs to promote organelle homeostasis ^25,33,34,81^. Additionally, our SEM analysis indicated that *Kif13b*^-/-^ cells exhibit more membrane bulges along the ciliary shaft compared to WT cells, which may reflect upregulated ciliary EV budding in the mutant cells. Thus, despite being localized to cilia and large EVs of mCCD cells, KIF13B appears to suppress release of large EVs that contain ciliary proteins. This is somewhat surprising, given that its homolog, KLP-6, was reported to promote EV shedding from the ciliary tip of male sensory neurons in *C. elegans*, with loss of KLP-6 instead causing excessive shedding of EVs into the glial lumen encompassing the ciliary base ^38^. Since it is currently unclear whether our *Kif13b*^-/-^ mCCD cells upregulate large EV shedding of ciliary proteins from the tip or base of cilia, it remains possible that large EVs containing PC2 are released from the ciliary base instead of the tip in these cells, similar to the scenario in *C. elegans klp-6* mutants. This would align with our observation that cilia of *Kif13b*^-/-^ mCCD cells contain increased membrane bulging along the side of cilia as compared to WT cells, although by SEM we only observe EVs that are just being budded and may therefore not be getting a representative view of how many side versus tip large EVs are actually being shed.

**Figure 7.**
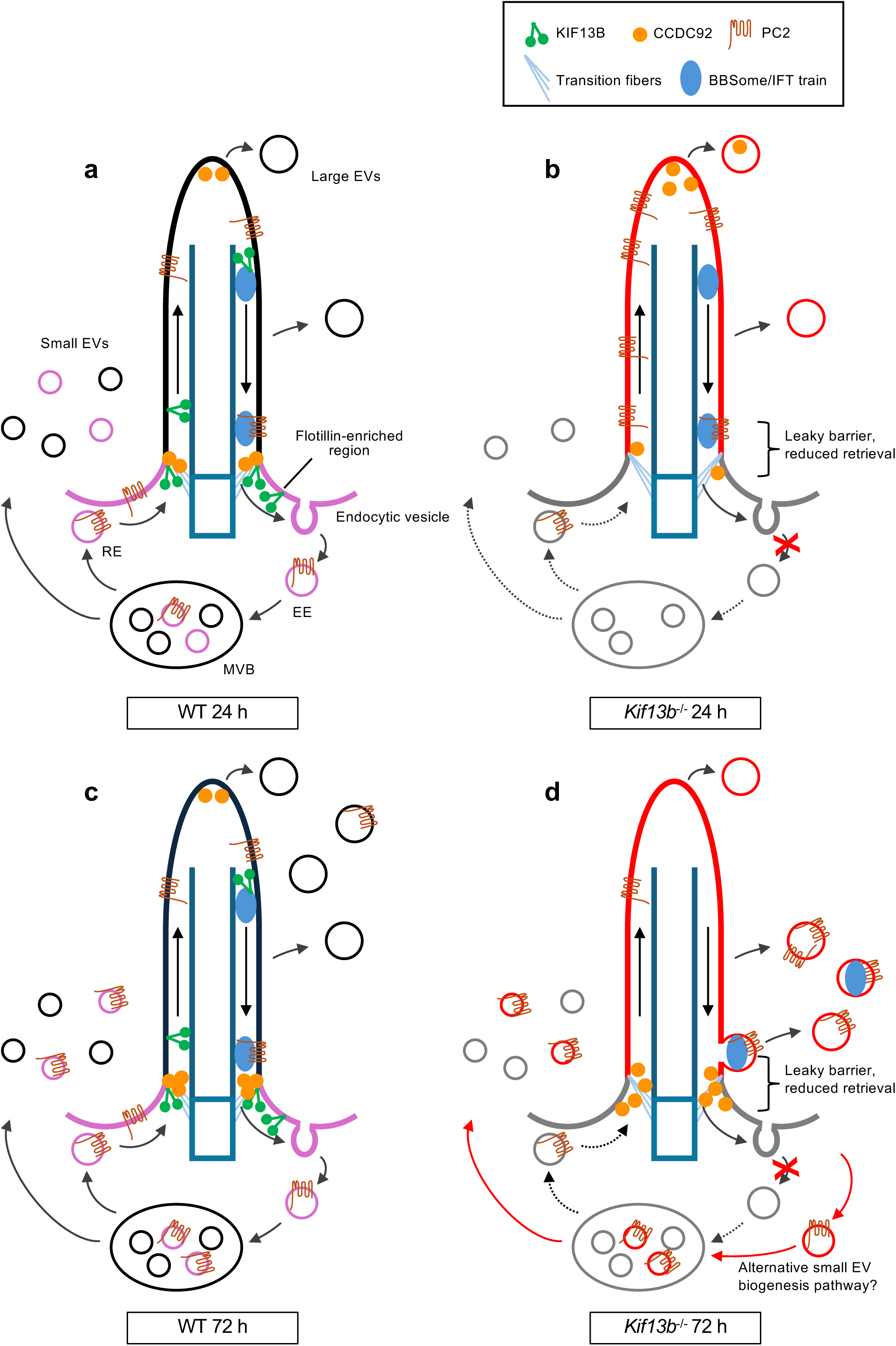
Summary of results and working model for how KIF13B affects ciliary PC2 homeostasis and EV release. (**a**) In WT cells starved for 24 h, KIF13B localizes dynamically along the cilium-basal body axis and is transported as retrograde IFT cargo from the ciliary tip to base ^44^. PC2 is not released into large or small EVs under these conditions, and CCDC92 localizes to the basal body transition fibers and ciliary tip. (**b)** 24 h serum starved *Kif13b*^-/-^ cells display increased ciliary levels of PC2 (and HGS and FLOT1; not shown), reduced release of small EVs generated by the ceramide/nSMase2-dependent pathway and altered composition of both small and large EVs. This is likely due to a combination of a leaky diffusion barrier and impaired endocytic retrieval at the ciliary base, which correlates with CCDC92 redistribution from the ciliary base to tip where it is released in large EVs. (**c**, **d**) After 72 h of serum starvation, WT cells release PC2 in both large and small EVs to control ciliary PC2 homeostasis. Large EV release of PC2 and other ciliary components is increased in *Kif13b*^-/-^ cells, leading to PC2 depletion from the mutant cilia, and CCDC92 is lost from the ciliary tip due to its release in large EVs and/or perturbed trafficking from the base to the tip. The *Kif13b*^-/-^ cells also upregulate the release of small EVs to WT levels, but mutant small EVs are devoid of cilia-derived proteins such as ZDHHC5 and are generated by an alternative pathway insensitive to nSMase2 inhibition.

While numerous ciliary and non-ciliary proteins were altered in abundance in large and small EVs isolated from *Kif13b*^-/-^ cells, reflecting the multiple cellular functions of KIF13B ^82,83^, it is noteworthy that two out of three Tier1 proteins that accumulated aberrantly within mutant large EVs at the 24 h time point, namely CCDC92 and CCDC198, localize to cilia. CCDC92 is a known centriole DAP that directly binds to the DAP CEP164 ^60^ and is phosphorylated by another DAP, TTBK2 ^62^. A recent study showed that TTBK2 additionally interacts with CSNK2A1, and when *Csnk2a1* was knocked out in mouse embryonic fibroblasts, cilia from mutant cells displayed aberrant protein content and accelerated EV release reminiscent of the phenotypes observed for our *Kif13b*^-/-^ mCCD cells ^63^. Intriguingly, the authors also found that EVs from *Csnk2a1* mutant cells were enriched for FLOT1, while FLOT1, FLOT2 as well as CAV1 were identified as proximity interactors of TTBK2 ^63^. This supports the notion that centriole DAPs may anchor to a specialized region of the periciliary membrane enriched for Flotillins (and CAV1), and contribute to regulating EV biogenesis or endosomal trafficking associated with this region. Given that KIF13B functions as a scaffold protein to regulate CAV1-mediated endocytosis ^84^ and CAV1 ciliary base localization ^43^, KIF13B could modulate ciliary protein content at the level of the transition fibers by regulating the interaction between DAPs and this specialized domain of the periciliary membrane (Fig. 7). Although it remains unclear if and how KIF13B interacts physically with CCDC92 or other DAPs, we note that CSNK2A1 was identified as a putative KIF13B interactor in our previously published KIF13B interactome analysis in human embryonic kidney (HEK) 293T cells ^85^, raising the intriguing possibility that KIF13B may affect DAP function via CSNK2A1. More work will be required to explore this possibility. Furthermore, while CCDC92 has been linked to cilia and DAPs in previous studies ^60,62^, its function at the cilium is not well understood. However, as several reports have linked CCDC92 to metabolic disorders such as obesity and diabetes ^86–88^, it will be interesting to investigate if and how these phenotypes are linked to defects in ciliary function and/or EV release.

In contrast to CCDC92, CCDC198 has not previously been shown to localize to the cilium or basal body, but a study found that loss of CCDC198 in the mouse led to altered body weight and decreased energy expenditure, which correlated with genome-wide studies in humans linking CCDC198 to higher body mass index ^64^, a common phenotype in ciliopathies ^2,3^. Notably, CCDC198 was reported to localize to the membrane of kidney epithelial cells and interacted directly with FNBP1L in a yeast two-hybrid screen ^64^. Since FNBP1L is a CDC42 effector that promotes actin-based endocytosis downstream of CDC42 ^65^, and activation of CCDC42 was shown to trigger release of EVs from cilia ^29^, it is tempting to speculate that KIF13B may promote release of CCDC198-containing EVs from cilia by a mechanism involving FNBP1L and CDC42, e.g. by upregulating ciliary CDC42 activity as seen by disruption of the BBSome ^29^. On the other hand, we were unable to detect significant alterations in ciliary CCDC198 localization in our *Kif13b*^-/-^ cells as compared to WT cells, hence more work is needed to investigate this further.

In addition to causing profound enrichment of ciliary proteins in large EVs after 72 h of starvation, loss of KIF13B also had a substantial impact on the small EV protein content at this time point. Specifically, the mutant small EVs lacked several proteins that were reported to be enriched in small EVs from BBSome-deficient KM cells ^20^, including, for example, the palmitoyl transferase ZDHHC5, which we found localizes to primary cilia and regulates ciliary length and PC2 levels. Although ciliary levels of ZDHHC5 were largely unaffected by loss of KIF13B, we found that ZDHHC5 accumulated aberrantly within cilia of *Ift27*^-/-^ IMCD3 cells as well as *Bbs4^-/-^*and *Bbs6^-/-^* KM cells, supporting that the BBSome mediates retrograde transport of ZDHHC5 out of cilia and that KIF13B could function downstream of the BBSome to promote its subsequent endocytic trafficking and small EV release. In support of this notion, we observed that the core BBSome component TTC8/BBS8 was also depleted from small EVs of 72 h serum-starved KIF13B mutant cells. Moreover, although the mechanism by which KIF13B regulates ZDHHC5 trafficking remains to be clarified, it is notable that ZDHHC5 binds to DLG4 ^89^, a direct interactor of KIF13B ^90^, suggesting that KIF13B may affect ZDHHC5 trafficking via DLG4 or its close homolog DLG1. Supportively, we have recently described an important role for DLG1 in regulating ciliary protein trafficking in mouse kidney epithelial cells, and showed that loss of DLG1 causes depletion of PC2 from cilia in mCCD cells ^48^, similar to our observations for *Zdhhc5*^-/-^ cells.

ZDHHC5 is a relatively well-studied and widely expressed protein that amongst other processes regulates synaptic plasticity, cardiac function, cell adhesion and fatty acid uptake ^91^. It has not previously been described to function at the primary cilium, but global knock out of *Zdhhc5* in the mouse was reported to cause a range of phenotypes, including a short snout, reduced retinal thickness, reduced fat tissue content, cardiac defects and unilateral hydronephrosis ^92^. Since several of the tissues and organs affected in *Zdhhc5* mutant mice are also affected in ciliopathies ^2,3^, it will be interesting to investigate if some of these phenotypes are linked to ciliary dysfunction. Moreover, as multiple ciliary proteins are known to be palmitoylated, but the responsible enzyme(s) has not yet been identified ^93^, future efforts to identify possible novel ciliary substrates of ZDHHC5 are warranted.

Finally, although the molecular basis for the upregulated rate of small EV release frequency observed in the *Kif13b*^-/-^ cells after 72 h of serum starvation remains to be clarified, it is notable that previous studies reported that mutations in genes that affect certain endo-lysosomal- or EV biogenesis pathways can lead to upregulation of alternative pathways to maintain cellular homeostasis. For example, in *Drosophila* motor neurons, opposing pathways involving RAB4, RAB11 and the retromer complex ensure plasticity of endo-lysosomal- and EV trafficking to maintain cellular homeostasis when key components of one pathway are mutated ^94^. Unraveling the mechanisms underlying such plasticity will be an important task for future studies and may help explain specific aspects of ADPKD disease pathology as well as other ciliopathies.

## Materials and Methods

### Cell lines and reagents

Cell lines and reagents used in this study are listed in Key Resources Table. Immortalized mCCD were generously provided by Dr. Eric Féraille from University of Lausanne, Switzerland, and have been described previously ^47^. Human embryonic kidney (HEK) 293T cells were from ATCC (cat. #CRL-3216) whereas previously described WT and *Arl6*^-/-^ and *Ift27*^-/-^ IMCD3 Flp-In cell lines were kindly supplied by Dr. Maxence Nachury from the University of California, San Francisco (UCSF), USA ^76^.

### Cell culture conditions

mCCD cells were grown at 37 °C with 5% humidified CO_2_ in DMEM/F12 with GlutaMaX (Thermo Fisher Scientific, cat. #10565018) supplemented with 2% fetal bovine serum (FBS, heat inactivated, Merck), 5 µg/ml insulin (Merck, cat. # I6634), 5 µg/ml holo-transferrin (Merck, cat. #T0665), 5 ng/ml EGF (Merck, cat. #SRP3196), 50 nM dexamethasone (Merck, cat. #D4902), 60 nM sodium selenite (Merck, cat. #S5261), 1 nM 3,3′,5-Triiodo-L-thyronine sodium salt (Merck, cat. #T6397), and 1% penicillin-streptomycin (Merck, cat. #P0781). To promote ciliation, mCCD cells were grown to ca. 80% confluency and then cultured at 37 °C with 5% humidified CO_2_ for 24, 48 or 72 h in DMEM/F12 – GlutaMaX supplemented with 60 nM sodium selenite and 5 µg/ml holo-transferrin. HEK293T and IMCD3 cells were cultured as described previously ^48^.

### Generation of knockout and transgenic kidney epithelial cell lines

To knock out *Kif13b* in mCCD cells we used CRISPR/Cas9 methodology essentially as described in ^48^. For *Kif13b* knock out, we utilized four sgRNA sequences from the mouse CRISPR “Brie” Knockout Library ^95^. The specific sequences are provided in Key Resources Table. After transfecting the cells with the generated Cas9-sgRNA plasmids and applying antibiotic selection, KIF13B protein depletion was assessed by western blot analysis. To knock out *Zdhhc5* Mammalian CRISPR Vector (Dual gRNA) was customized from VectorBuilder using the function “Design My Vector” and purchased. The plasmid codes for a Cas9-GFP fusion protein and two sgRNA against *Zdhhc5* exon 3, from the VectorBuilder database. The specific sequences are provided in Key Resources Table. Briefly, mCCD cells were transfected with the plasmid using Lipofectamine 3000 (Thermo Fisher Scientific). A day after transfection the GFP positive cells were FACS sorted into single cells in a 96 well plate with a FACS Aria III instrument. When cells reached confluency, they were sub-cultured in two 96 well plates. One plate was used to screen the clones for ZDHHC5 depletion by western blot analysis, the other plate was used to keep the clones in culture. Three knock-out clones were expanded and sent for Sanger sequencing at Eurofins Genomics. The clone that we used for this study has a deletion of 31 bp in exon 3 leading to a frameshift.

To create transgenic cell lines for live-cell imaging and/or rescue experiments, we generated lentiviral expression constructs coding for mNG-KIF13B, mCherry-KIF13B or mNG-ZDHHC5, expressed under EF1a and IRES promoters to ensure low and stable expression ^96^. In summary, the full-length human *KIF13B* or mouse *Zdhhc5* coding sequence was cloned into pENTR-mCherry-C1 or pENTR-mNG-C1 using standard cloning techniques. Primers utilized for this process are detailed in Key Resources Table. The resulting Gateway entry plasmids were further recombined with the pCDH-EF1a-Gateway-IRES-BLAST destination plasmid using the Gateway LR recombination kit (Invitrogen, cat. #11791020). All cloning vectors were kindly supplied by Dr. Kay Oliver Schink, Oslo University Hospital, Norway, and were described in ^97^. To generate lentiviruses, lentiviral expression plasmids were co-transfected with packaging plasmids pMD2.G and pCMVΔ-R8.2 (kind gift of Dr. Carlo Rivolta, Institute of Molecular and Clinical Ophthalmology Basel, Switzerland) into HEK293T cells using Lipofectamine 3000 (Invitrogen, cat. #L3000015). The clarified culture medium containing the lentivirus particles was subsequently used to transduce IMCD3 WT, IMCD3 *Ift27^-/-^*, mCCD WT, and mCCD *Kif13b*^-/-^ cells. Expressing cells were selected using blasticidin at a concentration of 5-15 µg/ml, and successful expression was confirmed through western blotting and IFM.

### SDS-PAGE and western blot analysis

Unless otherwise indicated, SDS-PAGE and western blotting were performed as described previously ^98^ by using antibodies and dilutions as listed in Key Resources Table. For western blot analysis of small EVs purified by ultracentrifugation, small EV pellets were resuspended in loading buffer (0.125 M Tris-HCl, 20% (v/v) glycerol, 0.004% (w/v) bromphenol blue) run on a 10% SDS-PAGE and transferred to a PVDF membrane (Immobilon® -FL PVDF membrane, Sigma-Aldrich, cat. #05317). The membrane was blocked with blocking buffer (0.5% Casein; 0.1% Tween; 0.04% NaN_3_; 150 mM NaCl) and incubated with primary (overnight, 4°C) and secondary antibodies (1 h), listed in Key Resources Table. Fluorescence detection was carried out using Odyssey Fc Imaging System (LICOR Bioscience).

### Immunofluorescence microscopy

Kidney epithelial cells were grown to ca. 80% confluency and then serum- and hormone-deprived for 24 or 72 h. Unless otherwise indicated, cells were first fixed with 4% PFA for 15 min at room temperature or 20 min at 4 °C followed by permeabilization with 1% BSA and 0.2 % Triton X-100 in PBS for 12 min. Permeabilized cells were blocked in 2% BSA blocking buffer for 1 h at room temperature. Cells were then incubated 1-2 h at room temperature or overnight at 4 °C with primary antibodies diluted in blocking buffer. KM cells were permeabilized in PBS with 0.3% Triton-X and for blocking and staining, fishblock solution with 0.3% Triton-X 100 was used. After several washes in PBS, cells were next incubated with secondary antibodies diluted in 2% BSA in PBS for 1 h at room temperature. Finally, nuclei were stained with DAPI (Sigma-Aldrich, cat. #D9542). Coverslips were mounted on slides with Shandon Immu-Mount (Thermo Scientific, cat. #9990402) supplemented with 0.5% N-propyl gallate.

For PC2 staining, we used an IFM protocol method described in ^49^ where we pre-fixed cells with 0.4% PFA for 5 min at 37 °C, permeabilized with PHEM buffer (50 mM PIPES; 50 mM HEPES; 10 mM MgCl_2_; pH 6.9) containing 0.5% Triton X-100 for 5 min at 37 °C, followed by a 4% PFA fixation for 15 min at room temperature. For HGS staining, we followed an IFM protocol method described in ^99^ where we briefly washed the cells with cytoskeletal buffer, then immediately fixed them with ice-cold MeOH inside a −20 °C freezer for 10 min. Imaging of IFM samples was done as described previously ^48^.

### Time lapse live cell imaging

To observe KIF13B and ZDHHC5 dynamics in the primary cilia of kidney epithelial cells, we used mCCD cells stably expressing mNG-KIF13B, and IMCD3 and mCCD cells stably expressing mNG-ZDHHC5. The primary cilia were visualized using SiR-tubulin staining. Cells were cultured on a 35 mm glass-bottom dishes and serum-starved for 24-72 h. For staining with SiR-tubulin dye (Cytoskeleton, Inc., cat. #CY-SC002), culture medium was supplemented with 1 µM of SiR-tubulin and cultures incubated in a cell incubator at 37 °C in a humidified atmosphere containing 5% CO_2_ for at least 30 min prior to cell imaging. Imaging was performed in a humidity chamber on an Olympus inverted microscope (IX83) equipped with Yokogawa spinning disc with 100× oil objective. The 488 nm and 640 nm laser lines were used for imaging mNG and SiR-Tubulin-Cy5, respectively. mCCD cells stably expressing mNG-KIF13B, which were serum starved for 24 h, were imaged using the Hamamatsu ORCA-Flash 4.0 digital camera (C13440) using the confocal settings at 5-10 sec time intervals. The rest of the cell lines and serum-starved conditions were imaged with the Teledyne Prime 95B back-illuminated sCMOS camera (01-PRIME-95B-R-M-16-C) using confocal settings at 3 sec time intervals.

### Purification of extracellular vesicles

Large EVs for MS analysis were purified by differential centrifugation of spent medium from 24 or 72 h serum-starved mCCD cells. To remove cell debris, the medium was first centrifuged at 2,000 x *g* for 20 min. This higher-speed initial centrifugation was chosen to improve the purity of the final pellet by reducing contamination from smaller debris. The supernatant, containing large EVs, was carefully transferred without disturbing the pellet. Large EVs and particles were then collected by centrifugation at 10,000 x *g* for 50 min at 4 °C and resuspended in 6 M Urea-Tris buffer (0.4 M Tris pH 7.8; 6 M urea) for further MS analysis. To purify small EVs for NTA, TEM and MS analysis we used a previously published ultracentrifugation approach ^20^. Briefly, spent medium was collected from 24, 48 or 72 h serum-starved mCCD cultures and first centrifuged at 1,000 x *g* for 10 min to remove cell debris. The supernatant was collected and subjected to further centrifugation at 10,000 x *g* for 30 min at 4 °C to sediment large EVs and particles. The supernatant was then subjected to ultracentrifugation at 100,000 x *g* for 2 h at 4 °C to sediment small EVs (Optima L-80K ultracentrifuge, Beckman Coulter, swing bucket SW28 rotor or Optima MAX-E ultracentrifuge, Beckmann Coulter, TLA-55 fixed-angle rotor). The final pellet was resuspended in particle-free PBS, Laemmli buffer or 6 M Urea-Tris buffer and used for downstream analyses.

To purify large and small EV for western blot analysis (Fig. S3b), we used a combined centrifugation and lectin-mediated precipitation approach ^59^. Spent medium from 72 h serum-deprived mCCD cultures was first centrifuged at 3,000 x *g* for 30 min to remove cell debris. The supernatant was then centrifuged for 40 min at 10,000 x *g* to sediment large EVs, which were collected, resuspended in PBS supplemented with protease inhibitor and centrifuged again for 60 min at 20,000 x *g* prior to SDS-PAGE and western blot analysis. Meanwhile, 1 μg/ml Concanavalin A from *Canavalia ensiformis* (Sigma-Aldrich, cat. # C2010) was added to the supernatant and the mixture put on slow stir for 16 h at 4 L. Subsequently, small EVs were collected by centrifugation for 60 min at 14,000 x *g*. The pellet was resuspended in PBS supplemented with protease inhibitor and centrifuged again for 60 min at 20,000 x *g* prior to SDS-PAGE and western blot analysis.

### Scanning electron microscopy (SEM)

To prepare the samples for SEM imaging, mCCD cells grown on glass coverslips and serum starved for 24 h or 72 h were fixed in 2% glutaraldehyde in 100 mM sodium cacodylate at room temperature for at least 2 h up to several days. Subsequently, cells were washed three times in 100 mM sodium cacodylate and then treated with 1% OsO_4_ in 100 mM sodium cacodylate for 1-2 h at room temperature. The cells were then washed with deionized water and dehydrated with a graded series of ethanol solutions (30%, 50%, 70%, 80%, 95%, and 100%) at room temperature, with each step lasting 5-10 min. The samples were allowed to dry using a critical point drier. Dried samples were mounted onto stubs and sputter coated with 6 nm gold. The coated samples were then placed into a SEM chamber and images captured with a FEI Quanta 3D FEG dual beam SEM microscope (University of Copenhageńs Core Facility for Integrated Microscopy).

### Transmission electron microscopy (TEM) of small EVs

Small EV pellets isolated by ultracentrifugation from spent media of mCCD cells following 48 h serum-starvation were resuspended in particle-free PBS. They were subsequently applied to 400 mesh carbon-coated copper grids (Electron Microscopy Science) and allowed to settle for 1 min. Embedded EVs were fixed with 1% glutaraldehyde for 5 min and washed four times with deionized water (ddH_2_O). Negative staining was performed by applying 2% uranyl acetate for 1 min and the grids were allowed to dry. Imaging was done with a FEI Tecnai G2 12 BioTwin transmission electron microscope (Fei, Company, Hillsboro, USA).

### Nanoparticle tracking analysis (NTA)

Small EVs were isolated by ultracentrifugation from spent media of 24, 48 and 72 h serum-deprived mCCD cells. Inhibition of EV release was performed by treatment with 10 µM GW4869 (Sigma-Aldrich, cat. #D1629) dissolved in DMSO during serum starvation (24 h) prior EV preparation. Particles were resuspended in particle-free PBS and measured using ZetaView Nanoparticle Analyzer PMX-230 (TWIN) (Particle Metrix). Particle movement was recorded in scatter mode 1-2 Cycles at 11 Positions, at a steady temperature of 25°C and analysed using the Software ZetaView (version 8.05.16 SP3).

### Liquid chromatography (LC)-MS/MS analysis of EVs

Small EV pellets from spent medium of serum-deprived mCCD cells (n=4/cell line, 24 h and 72 h) were isolated using ultracentrifugation. Pellets were diluted in 6 M Urea-Tris buffer and subjected to liquid chromatography-MS analysis as described previously ^20^. For analysis of large EVs isolated by centrifugation, the protein samples were subjected to proteolytic cleavage as described earlier ^20^ before the resulting peptides were analysed by LC-MS. Peptide separation was performed using a nanoElute 2 LC system (Bruker) coupled to a timsTOF Ultra 2 mass spectrometer (Bruker). A trap-column chromatography setup was employed, consisting of a Thermo Trap Cartridge (5 mm) for sample trapping and a PepSep Twenty-five Series analytical column (75 µm ID, 1.5 µm particle size). The column temperature was maintained at 50 °C throughout the analysis.

Samples were loaded using an ultra-low volume (µl) pickup injection method, with a total sample loading volume of four pick-up cycles plus two microliters. Chromatographic separation was performed at a constant flow rate of 0.30 µl/min, using a gradient elution with mobile phase A consisting of 0.1% formic acid in water and mobile phase B of 0.1% FA in 99.9% acetonitrile. The gradient started at 5% B, increasing to 25% B at 24 min, then to 35% B at 30 min, and reaching 95% B at 34 min, where it was held until 38 min before re-equilibrating to 5% B at 40 min.

Data acquisition was performed in DIA-PASEF mode (Data-Independent Acquisition Parallel Accumulation–Serial Fragmentation) ^100^. Ionization was performed using a CaptiveSpray nanoelectrospray source, with a capillary voltage of 1600 V. The dry gas flow rate was set to 3.0 l/min, and the drying temperature was maintained at 200 °C. The instrument was operated in positive ion mode, with Trapped Ion Mobility Spectrometry (TIMS) enabled to enhance precursor separation and fragmentation efficiency. Spectra were acquired across a mass range of 100 to 1700 m/z, with an ion mobility range (IM) from 0.60 to 1.40 Vs/cm². The accumulation time per scan was 2.0 milliseconds, with a ramp time of 100.0 milliseconds. Each acquisition cycle included 12 DIA-PASEF scans, resulting in a total cycle time of approximately 1.38 seconds. DIA windows were dynamically adjusted based on precursor density, covering an m/z range of 300 to 1200. Fragmentation was conducted using collision-induced dissociation (CID), with collision energy dynamically adjusted as a function of ion mobility. High-sensitivity detection mode was enabled to maximize signal acquisition efficiency.

### MS data analysis

Data-independent acquisition (DIA) data were processed using DIA-NN (Data-Independent Acquisition by Neural Networks) ^101^ version 1.9.2. The analysis was performed using a reference spectral library generated in silico from the Swissprot Mouse FASTA database (version 12/2024) and common contaminants database included in DIA-NN and refined empirically from the acquired DIA data. Search settings included N-terminal methionine excision, cleavage specificity at lysine (K) and arginine (R) with one missed cleavage allowed, and cysteine carbamidomethylation as a fixed modification. Variable modifications were restricted to one per peptide. The precursor and fragment mass accuracies were set to 15 ppm, Match Between Runs (MBR) and peptidoform-level scoring were enabled. Protein inference was performed using heuristic protein grouping, reducing redundancy while preserving quantitative accuracy. False discovery rate (FDR) filtering was applied at 1% at the precursor and protein levels.

Spectral library generation and retention time alignment were performed during a first-pass search, followed by reanalysis using the empirically refined library. Quantification was carried out at the precursor, protein, and gene levels. The “PG.MaxLFQ” expression matrix output reports normalized quantity employing the MaxLFQ principle ^102^ was used for quantitative analysis on the protein level for downstream statistical analysis. Ratio, t-test and significance A were calculated using the Perseus framework ^103^. Only proteins identified and quantified in more than 50% of the samples were considered. Missing values were replaced before statistical testing. For both statistical tests, correction for multiple testing was performed (t-test: permutation-based FDR; Significance A: Benjamini-Hochberg FDR). Proteins, detected to be significant for both tests after FDR determination with an FDR of 0.05 were considered as Tier 1 hits. Proteins with a p-value < 0.05 for both tests without FDR determination were considered as Tier 2 hits.

### GO term enrichment analysis

Gene ontology (GO) term enrichment analysis was performed using the PANTHER Overrepresentation Test (version released 20240807) available at geneontology.org. The analysis was conducted on a gene list derived from *Mus musculus* using the complete *Mus musculus* gene database as the reference list. The GO annotation database used was DOI: 10.5281/zenodo.14083199, released on 2024-11-03. Statistical significance was assessed using Fisher’s exact test, and multiple testing correction was applied using the FDR method. We searched for GO term enrichment for the categories molecular function and cellular component.

### Quantitative and statistical analysis of IFM, western blot and NTA data

Unless specified otherwise, quantitative and statistical analysis of IFM data was done as described previously ^48^. IFM images of HGS and ZDHHC5 staining in IMCD3 cells, and of ciliary length and PC2 staining in WT and *Zdhhc5*^-/-^ mCCD cells, were analyzed by a semi-automated method using the CiliaQ plugin on ImageJ. Rapidly, for each stack, borders were cropped, and the image center was treated with the CiliaQ Preparator v0.1.2 plugin, to detect and generate masks of cilia. The following parameters were used: Segmentation style: Set intensities above thresholds to maximum possible intensity value (creates a binary image). Channel duplicated to include a copy of the channel that is not segmented. Subtract Background: 20. Segmentation method: Canny 3D Thresholder, (plugin by Sebastian Rassmann, see https://github.com/sRassmann/canny3d-thresholder for a descriptions). Gauss sigma: 1.0. Canny alpha: 5.0. Low threshold method (hysteresis thresholding): IsoData. High threshold method (hysteresis thresholding): Otsu. The results images were treated by the CiliaQ Editor v0.0.3 plugin and unwanted detected object were discarded. Finally, we used the CiliaQ v0.1.7 plugin to extract protein of interest fluorescence intensities inside cilia masks ^104^. For each cilium, integrated fluorescence intensities were normalized by the average fluorescence intensities inside the cilium. For each experiment, intensities were further normalized to relevant control/WT condition. Each dot corresponds to one cilium and plotted data on graph originate from three independent experiment.

For western blot analysis we used Fiji software ^105^ to measure the average pixel intensity of the bands. Subsequently, we normalized the acquired numbers to the loading control and performed the statistical analysis in GraphPad Prism 10. All quantitative data are presented as meanL±Lstandard deviation. Differences were considered significant when the p-value was <0.05. Unless otherwise stated, the results were confirmed in at least three independent biological replicates. Statistical analysis on NTA data was done in Graphpad Prism 7.0 software (Graphpad Software Inc.). Outliers were excluded using the ROUT method (Q = 1%). Gaussian distribution was tested using Shapiro-Wilk and Kolmogorov-Smirnov test (P < 0.05). For Gaussian distributed data, *t* test and for non-Gaussian distributed data, Mann-Whitney *U* test performed. Significance levels: P > 0.05 not significant (ns), P ≤ 0.05 *, P ≤ 0.01 **, P ≤ 0.001 ***, P ≤ 0.0001 ****.

## Supporting information

Supplemental Tabl 5

## Acknowledgements

We are grateful to Feng Qian and the Polycystic Kidney Disease Research Resource Consortium (U54DK126114) for providing the anti-PC2 (rabbit polyclonal) antibody, and to Eric Féraille, Maxence Nachury, Kay Oliver Schink, Carlo Rivolta and Yuko Mimori-Kiyosue for cell lines and other reagents. We thank Geyi Li, Liva Karlshøj Julegaard, Emma Rose Jensen, Bjørk Ingeborg Bartholdy and Jannis von Spreckelsen for assistance with quantitative analysis of microscopy data. We thank the staff of the Core Facility for Medical Proteomics, University of Tübingen for excellent support. This work was supported by grants NNF18SA0032928 and NNF22OC0080406 from the Novo Nordisk Foundation (LBP), grant 2032-00115B from the Independent Research Fund Denmark (LBP), the European Union’s Horizon 2020 research and innovation program Marie Sklodowska-Curie Innovative Training Networks (ITN) grant 861329 (LBP, STC, KB), the TheRaCil consortium funded by the European Union (Horizon-health-2022-disease-06-two stage, grant 101080717), and the Deutsche Forschungsgemeinschaft FOR5547 – Project-ID MA 6139/6-1 503306912 (HMS).

## Author contributions

CKR, AF, FC, AMF, ALWP, SLJ, JKTS, AS, JL, CRB, KB and ZA performed experiments; CKR, AF, FC, BM, AMF, ALWP, CRB, KB, MC, HMS and ZA analyzed data; CKR, AF, FC, BM, STC, KB, MC, ZA, HMS and LBP prepared the figures; MB supplied reagents; ZA, HMS and LBP conceived the project; LBP wrote the paper with input from all authors; ZA, STC, KB, HMS and LBP obtained funding for the study.

## Key Resources Table

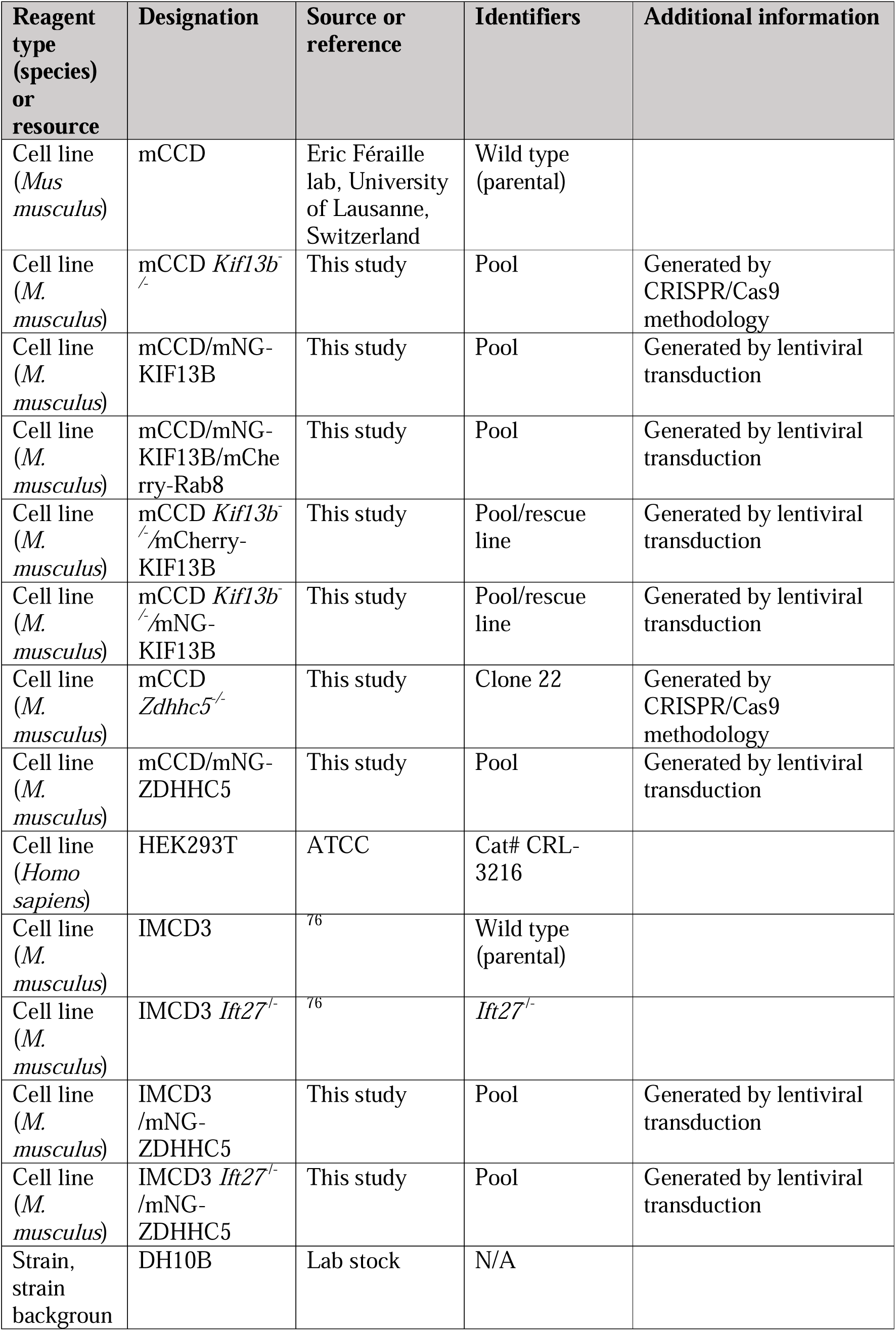
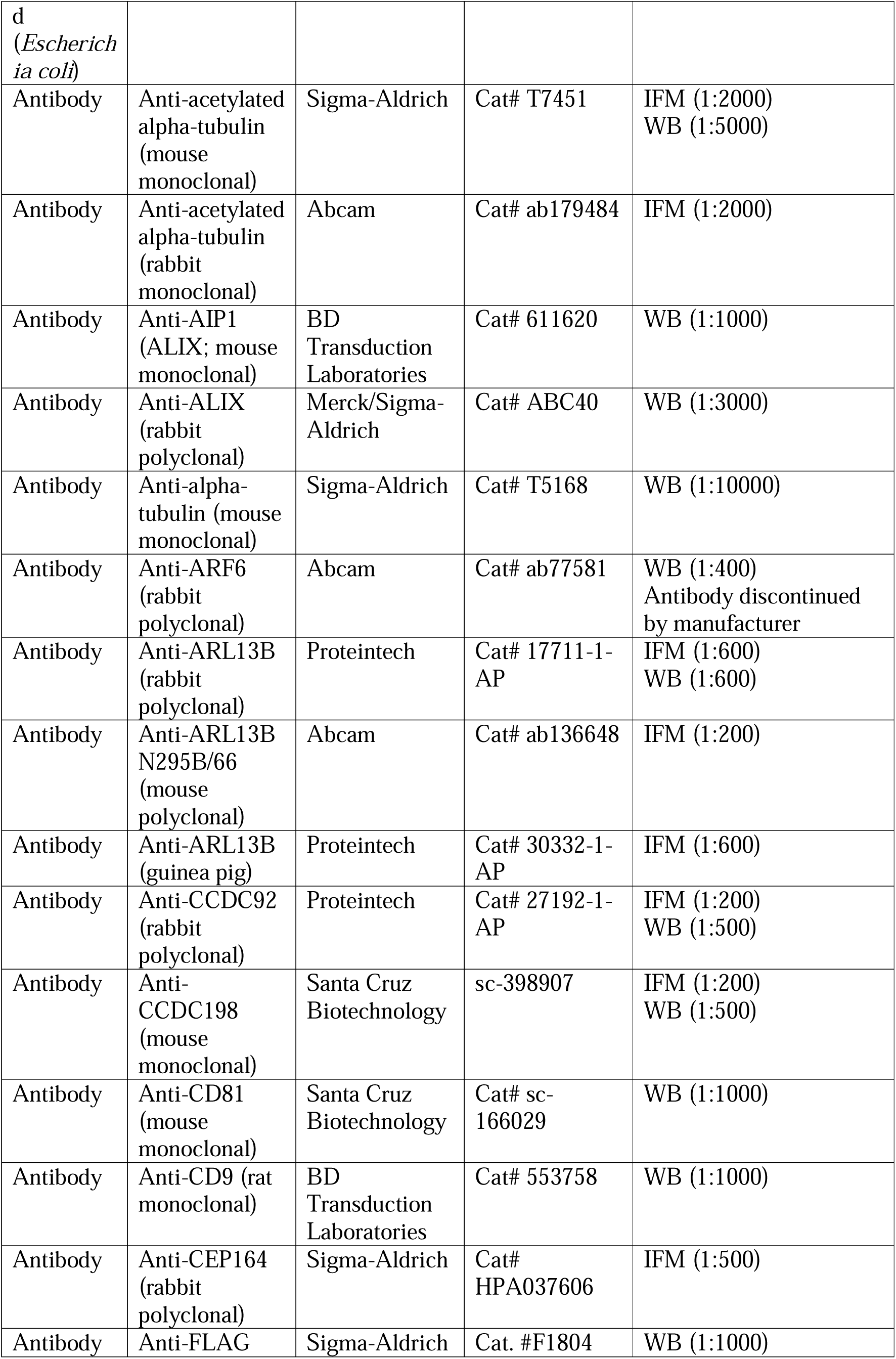
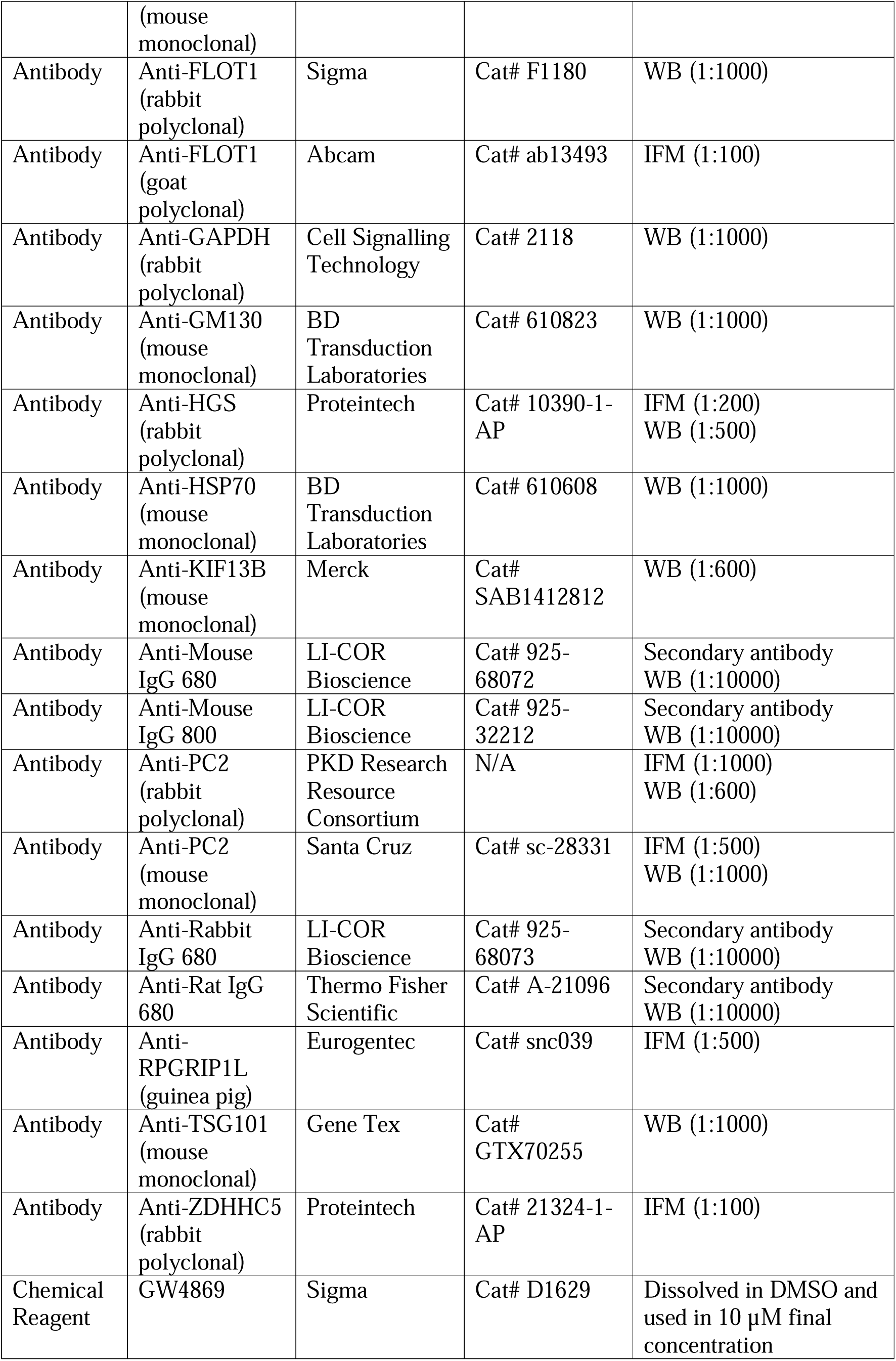
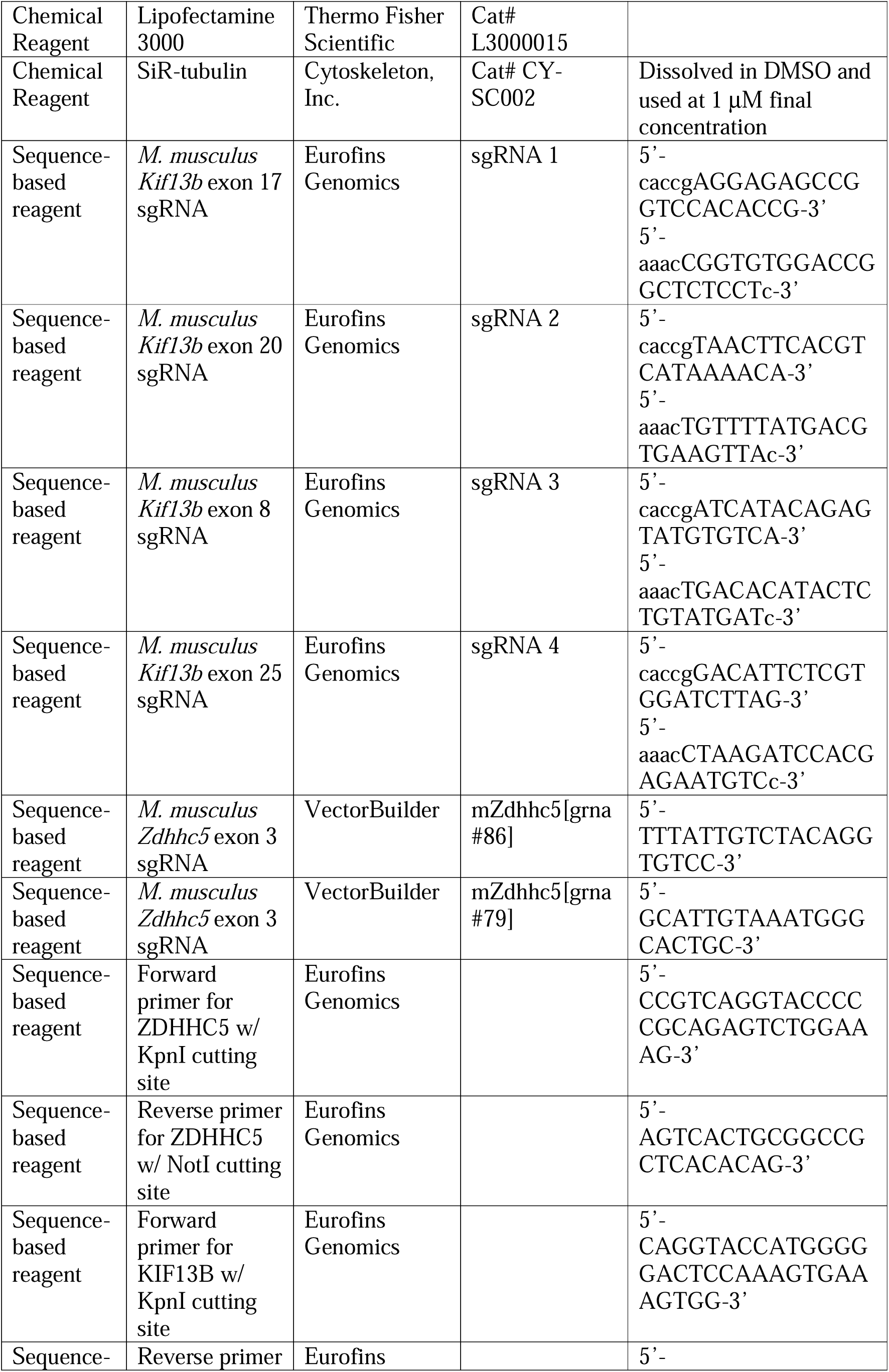
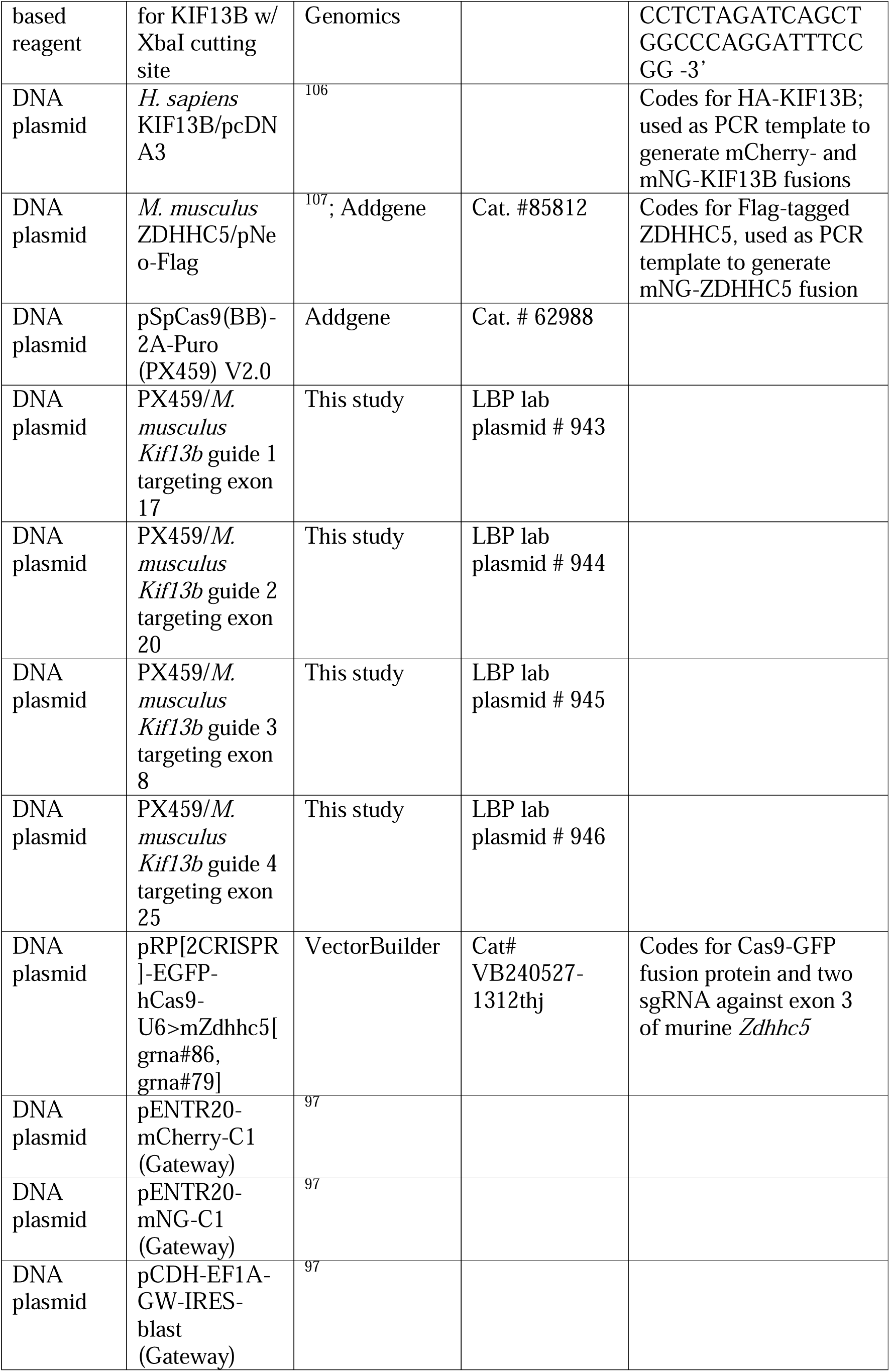
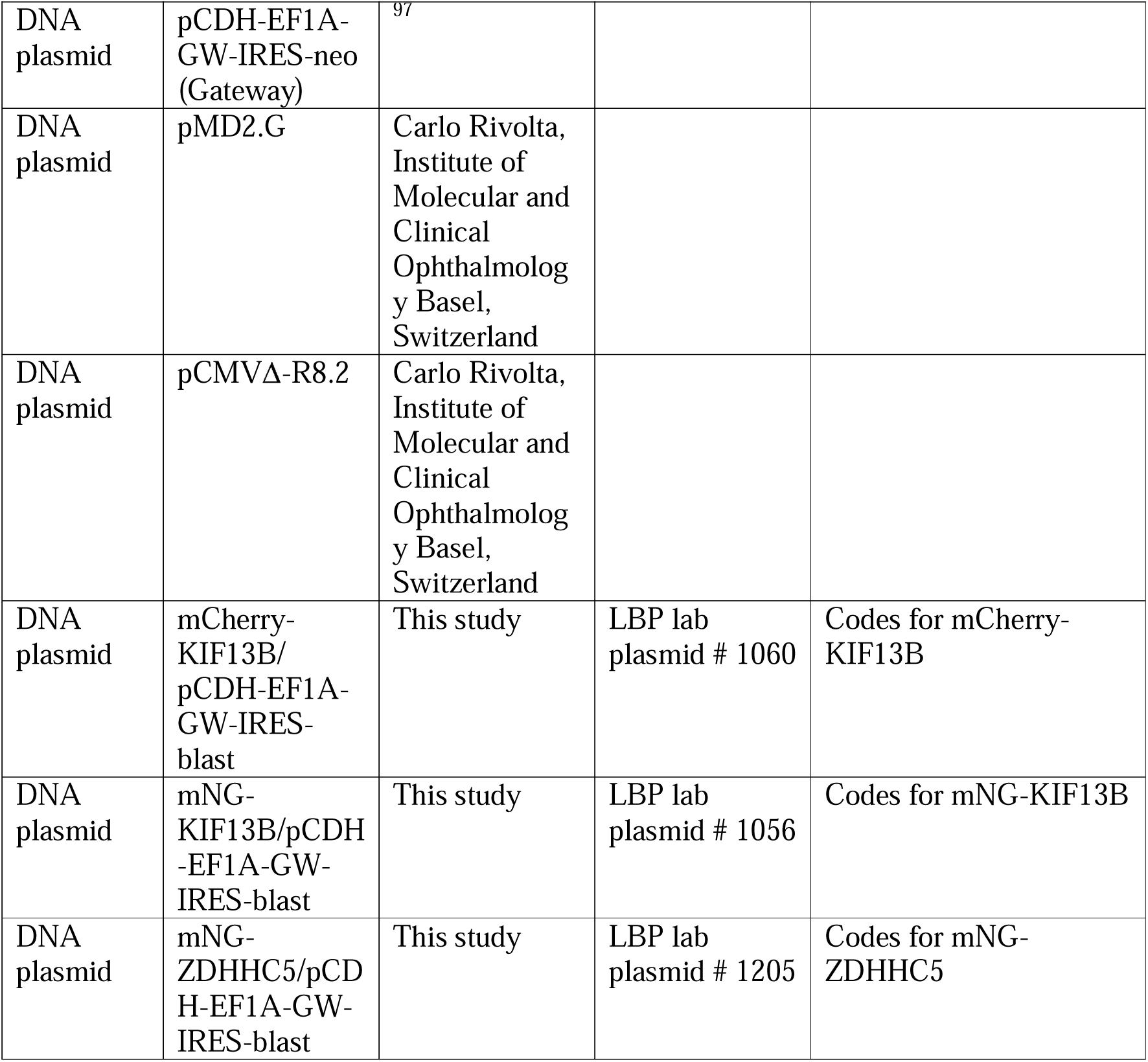

## Figure legends

**Movie 1. Burst-like intraciliary movement of mNG-KIF13B in live mCCD cells starved for 24-48 h.** Representative live cell imaging of *Kif13b*^-/-^ mCCD cells stably expressing mNG-KIF13B. Cells were serum starved for 24-48 h and stained with SiR-tubulin (magenta) to view the ciliary axoneme and microtubule cytoskeleton. Transient accumulation of mNG-KIF13B was observed in 15 out of 72 cilia (21%) imaged by this approach.

**Movie 2. Burst-like intraciliary movement and tip shedding of mNG-KIF13B in live mCCD cells starved for 72 h.** Representative live cell imaging of *Kif13b*^-/-^ mCCD cells stably expressing mNG-KIF13B. Cells were serum starved for 72 h and stained with SiR-tubulin (magenta) to view the ciliary axoneme and microtubule cytoskeleton. Transient accumulation of mNG-KIF13B was observed in 18 out of 57 cilia (32%) imaged by this approach.

**Movie 3. Ciliary localization of mNG-ZDHHC5 in mCCD cells.** Representative live cell imaging of WT mCCD cells stably expressing mNG-ZDHHC5. Cells were serum starved for 24 h and stained with SiR-tubulin (magenta) to view the ciliary axoneme and microtubule cytoskeleton.

**Figure S1. Characterization of WT, *Kif13b* mutant and rescue mCCD cell lines.** (**a**) Representative IFM images of 72 h serum-deprived WT and *Kif13b*^-/-^ cells, stained with anti-acetylated α-tubulin (AcTub) antibody to mark cilia. (**b**) Measurement of ciliation frequency in 24 or 72 h serum-deprived WT and *Kif13b*^-/-^ cells, based on IFM images as in (a). (**c**) Western blot analysis of lysates from the indicated mCCD cells lines subjected to serum starvation for 24, 48, or 72 h, as indicated, and probed with antibodies specific for the proteins listed on the right; molecular mass markers are shown in kDa to the left. (**d**) Western blots of indicated cell lines using antibodies against KIF13B and α-tubulin (α-tub; loading control). Molecular mass markers are shown in kDa to the left. mNG-KIF13B: mNG-KIF13B. (**e**) IFM analysis of *Kif13b*^-/-^ cells stably expressing mCherry-KIF13B or mNG-KIF13B after 24 h of serum starvation. The cells were stained with antibody against acetylated α-tubulin (AcTub). Cells expressing mCherry-KIF13B were additionally stained with an antibody against mCherry (red). Images to the right show enlarged regions of the boxed areas. Note that the KIF13B fusion proteins are concentrated at both centrioles at the base of primary cilia.

**Figure S2. Quantification of SEM data.** (**a**, **b**) Quantification of ciliary length (**a**) and percentage of cilia with a pocket (**b**) in indicated cell lines, based on SEM images as shown in Figure 2 (n=3). Between 61-67 cilia in total were measured for the 24 h samples (n=3), and 163-208 for the 72 h samples (n=4). *, p<0.5.

**Figure S3. Western blot analysis of small and large EVs from WT and *Kif13b*^-/-^ mCCD cell cultures.** (**a**) Scheme of procedure used for initial purification of large EVs and small EVs using centrifugation and lectin-mediated precipitation. (**b**) Large and small EVs were purified according to the scheme in (a) using spent medium of 72 h serum starved cultures; the EV samples were analyzed by western blotting using antibodies as indicated (AcTub, acetylated α-tubulin). Whole cell lysates are included for comparison, and molecular mass markers are indicated in kDa to the left. (**c**) Quantification of the relative distribution of the indicated proteins in large and small EVs, based on western blots as in (c), n=4. *, p<0.05; **, p<0.01; ns, non-significant. (**d**) Western blot analysis of small EVs purified from spent medium of 48 h serum starved cultures by ultracentrifugation (see Fig. 4a). Blots were probed with antibodies as indicated. Molecular mass markers are shown in kDa to the left. (**e**) Western blot analysis of whole cell lysates from the indicated cell lines, serum-starved for 24 or 72 h. Blots were probed with antibodies listed to the right, with DCTN1 serving as loading control; molecular mass markers are shown in kDa to the left.

**Figure S4. Ciliary localization of CCDC198**. (**a**) IFM analysis of indicated mCCD cell lines after 24 or 72 h of serum starvation, stained for ARL13B (magenta), CEP164 (red) and CCDC198 (green). mCh-KIF13B: mCherry-KIF13B. (**b**) Quantification of relative ciliary CCDC198 staining intensities of the indicated mCCD cell lines, based on images as shown in (a). ns, non-significant. (**c**) IFM analysis of the *Kif13b*^-/-^ mCCD cells stained with indicated antibodies. Note the co-localization of CCDC92 with acetylated tubulin (AcTub) in an EV-like particle near the ciliary tip (arrow).

**Figure S5. Ciliary localization of ZDHHC5 in WT and *Kif13b*^-/-^ or *Ift27*^-/-^ cell cultures.** (**a**, **b**) Representative IFM images (**a**) and quantification of relative ciliary levels of ZDHHC5 (**b**) in indicated mCCD cell lines serum-starved for 24 or 72 h and stained with antibodies against acetylated α-tubulin (AcTub; red) and ZDHHC5 (green). mCh-KIF13B, mCherry-KIF13B. Between 40-50 cilia were measured per condition per experiment (n=3) and the MFI of ciliary ZDHHC5 staining intensity normalized to the mean MFI for WT at 24 h. (**c**) WT and *Kif13b*^-/-^ mCCD cells stably expressing mNG-ZDHHC5 were serum starved, treated with SiR-tubulin to label cilia, and the percentage of cells with mNG-ZDHHC5 signal in cilia quantified based on 31 and 36 live cell imaging movies for the WT and mutant, respectively (see Movie 3 for representative image). (**d**) WT and *Ift27*^-/-^ IMCD3 cells stably expressing mNG-ZDHHC5 were treated as in (c), and the relative ciliary intensity of mNG-ZDHHC5 analyzed, based on 24 and 40 movies for the mutant and WT, respectively. **, p<0.01; ****, p<0.0001; ns, non-significant.

**Figure S6. Loss of ZDHHC5 affects ciliary length and PC2 levels in mCCD cells.** (**a**, **c**) Western blot analysis of WT and *Zdhhc5*^-/-^ cells, using antibodies as indicated. GAPDH serves as loading control; molecular mass markers are shown in kDa to the left. (**b**) IFM analysis of WT and *Zdhhc5*^-/-^ cells, using indicated antibodies. (**d**, **e**, **f**) Quantification of ciliation frequency (**d**), ciliary length (**e**), and relative ciliary PC2 intensity (**f**) of indicated cell lines serum-starved for 24 or 72 h. In (d), images of 40-50 cells were analysed manually per condition per experiment (n=2). For (e), we used CiliaQ ^104^ to automatically measure cilia length (n=2), and for (f), quantitative analysis was done by a semi-automated approach (see Materials and Methods) and normalized to the mean ciliary MFI of PC2 for WT (n=2). *, p<0.05; ***, p<0.001; ****, p<0.0001; ns, non-significant.

## Notes

### Competing Interest Statement

The authors have declared no competing interest.

### Summary of Updates

The manuscript has changed substantially, most figures have been updated, some have been deleted and new ones included. The author list has been changed slightly to reflect these changes.

