## Supplementary figures and images for "KIF13B controls ciliary protein content by promoting endocytic retrieval and suppressing release of large extracellular vesicles from cilia"

### Supplemental Figures

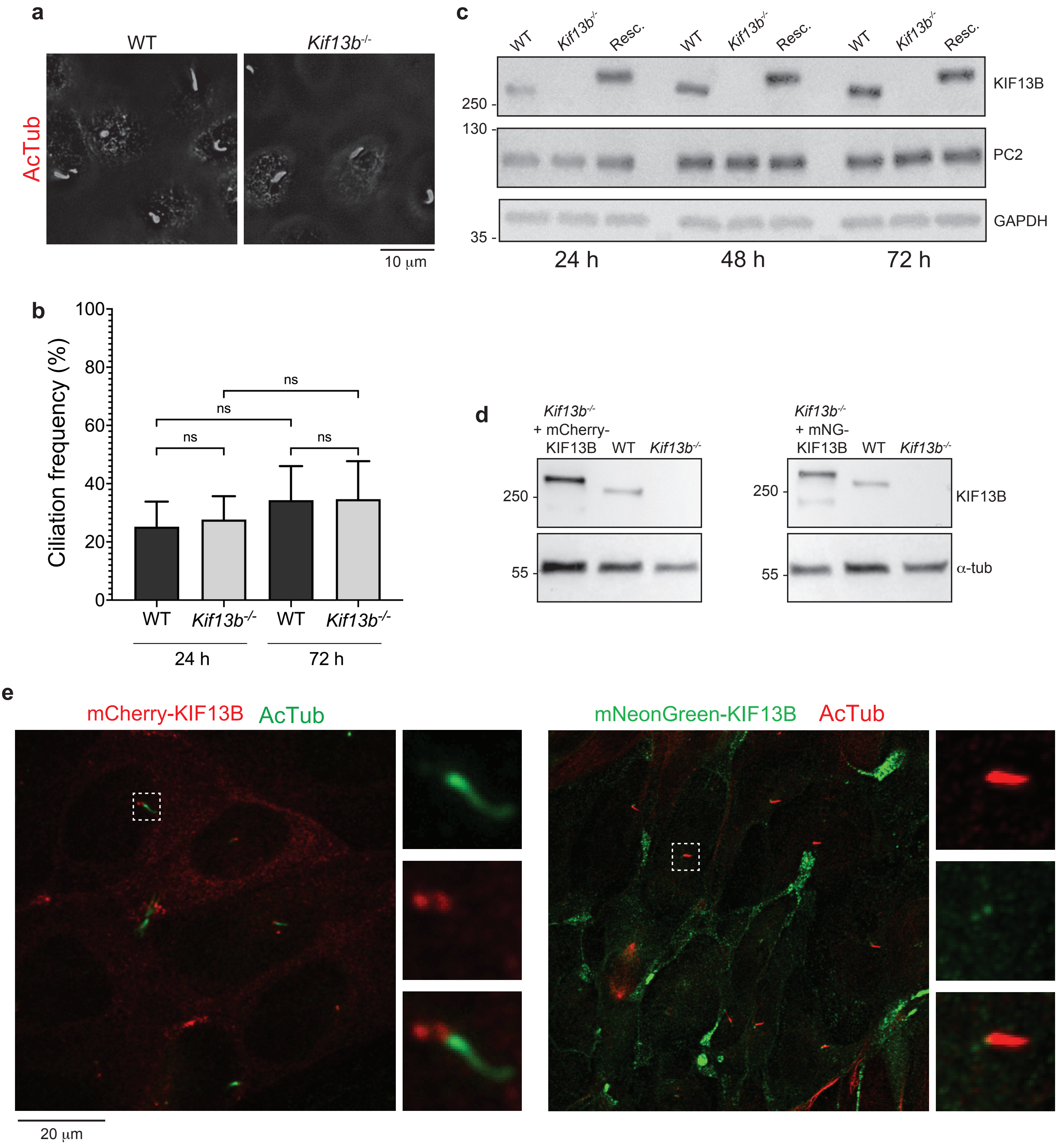

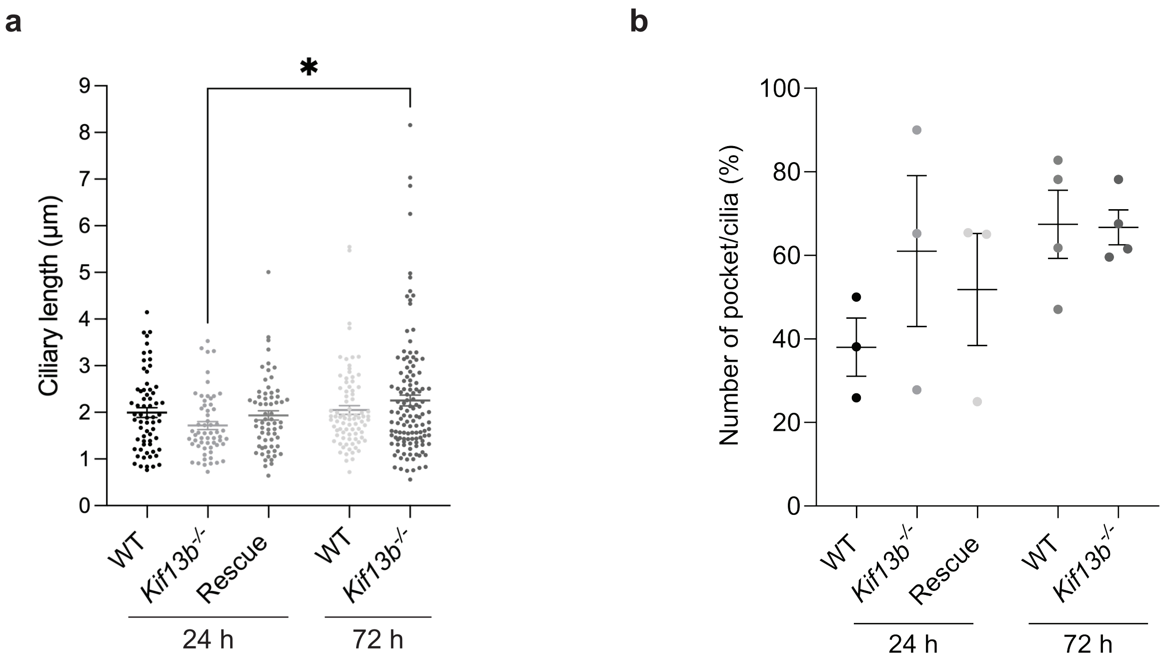

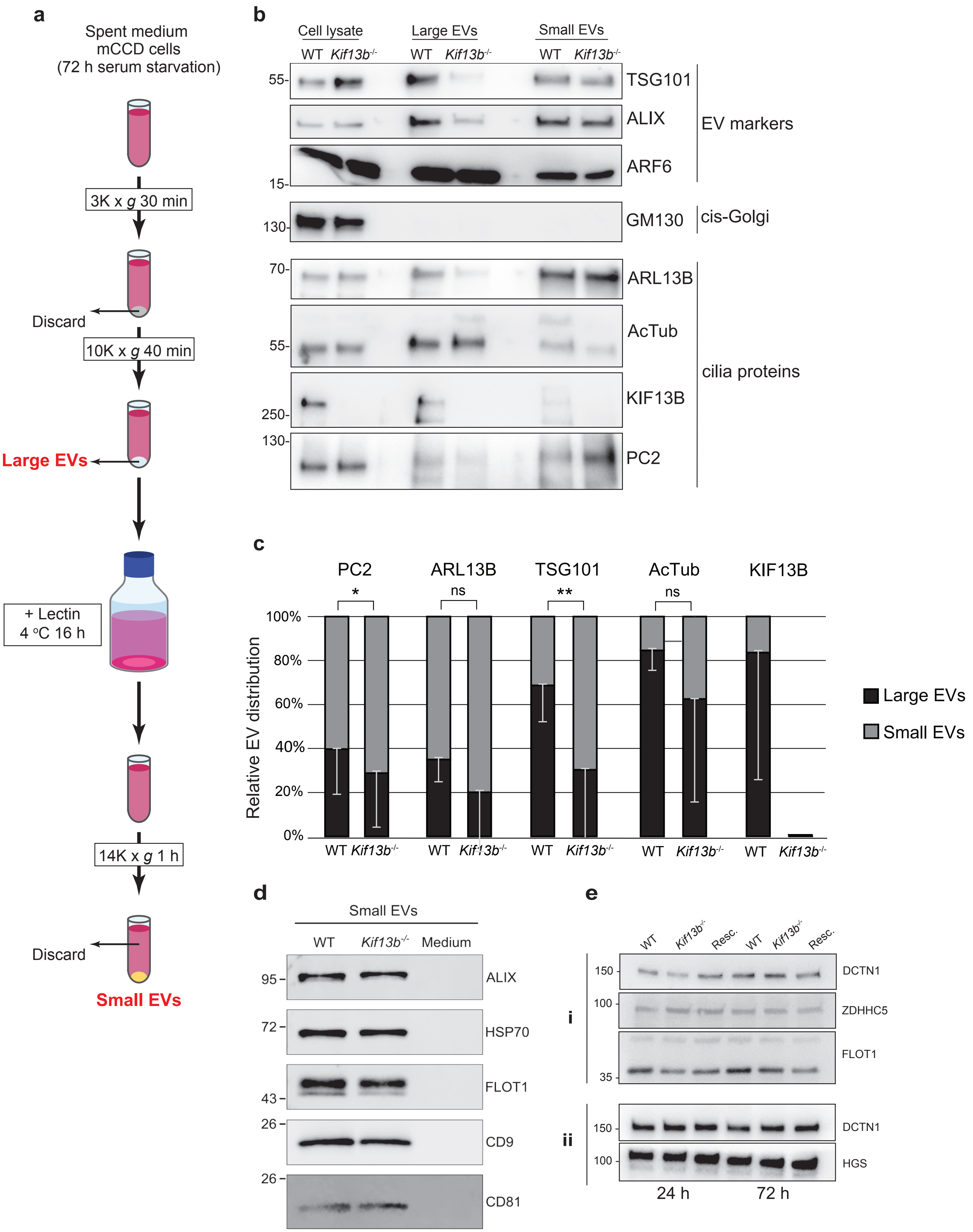

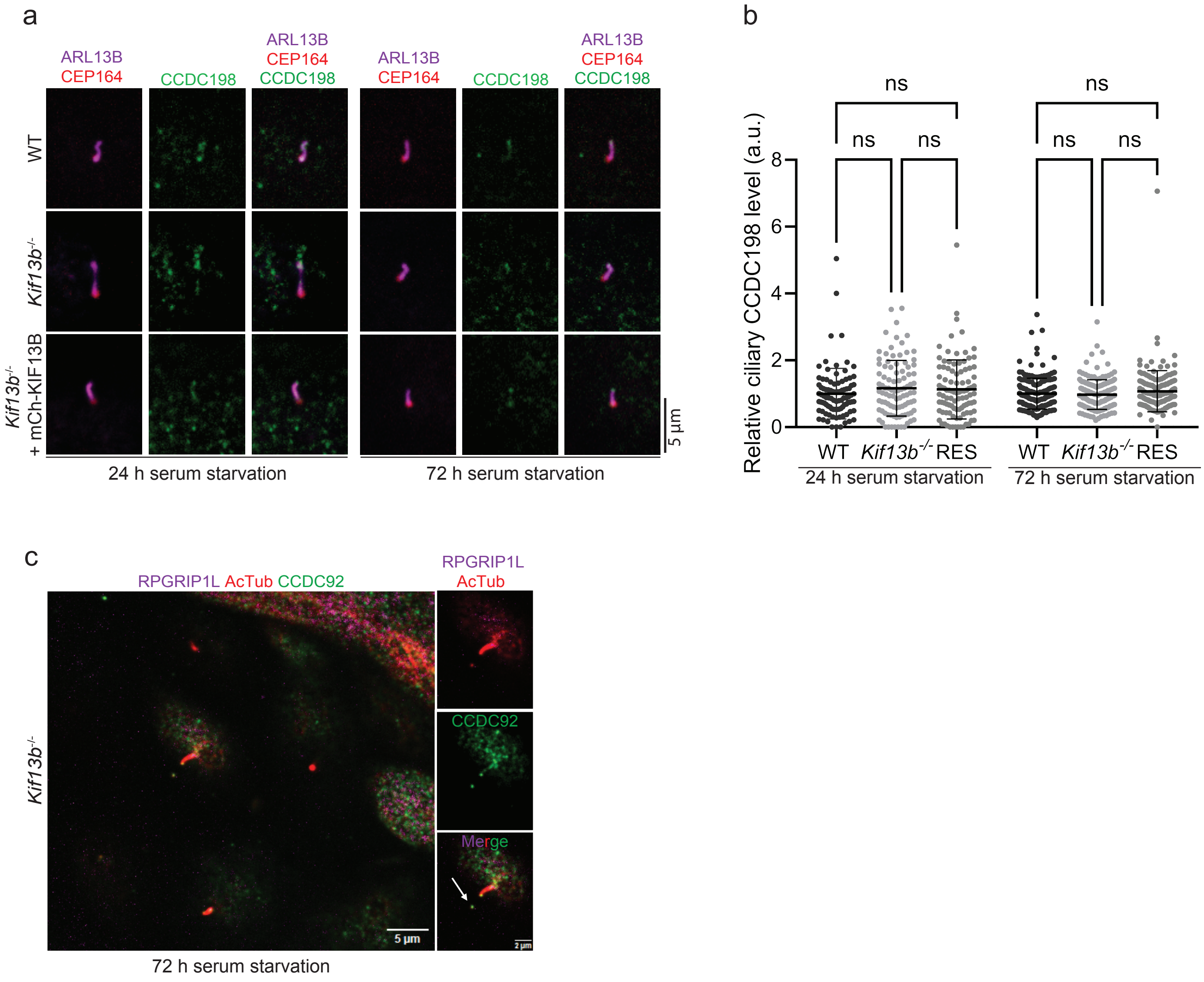

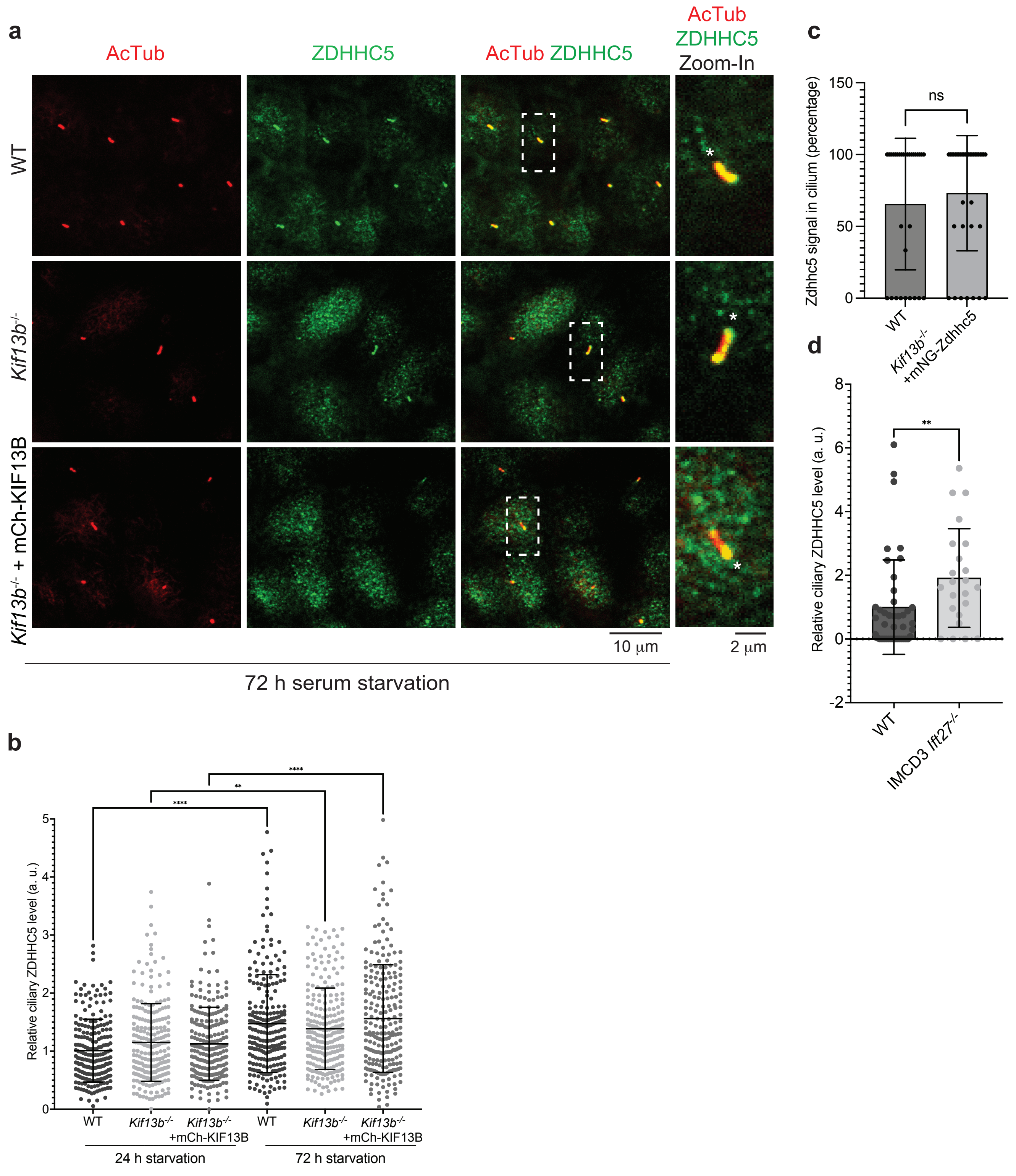

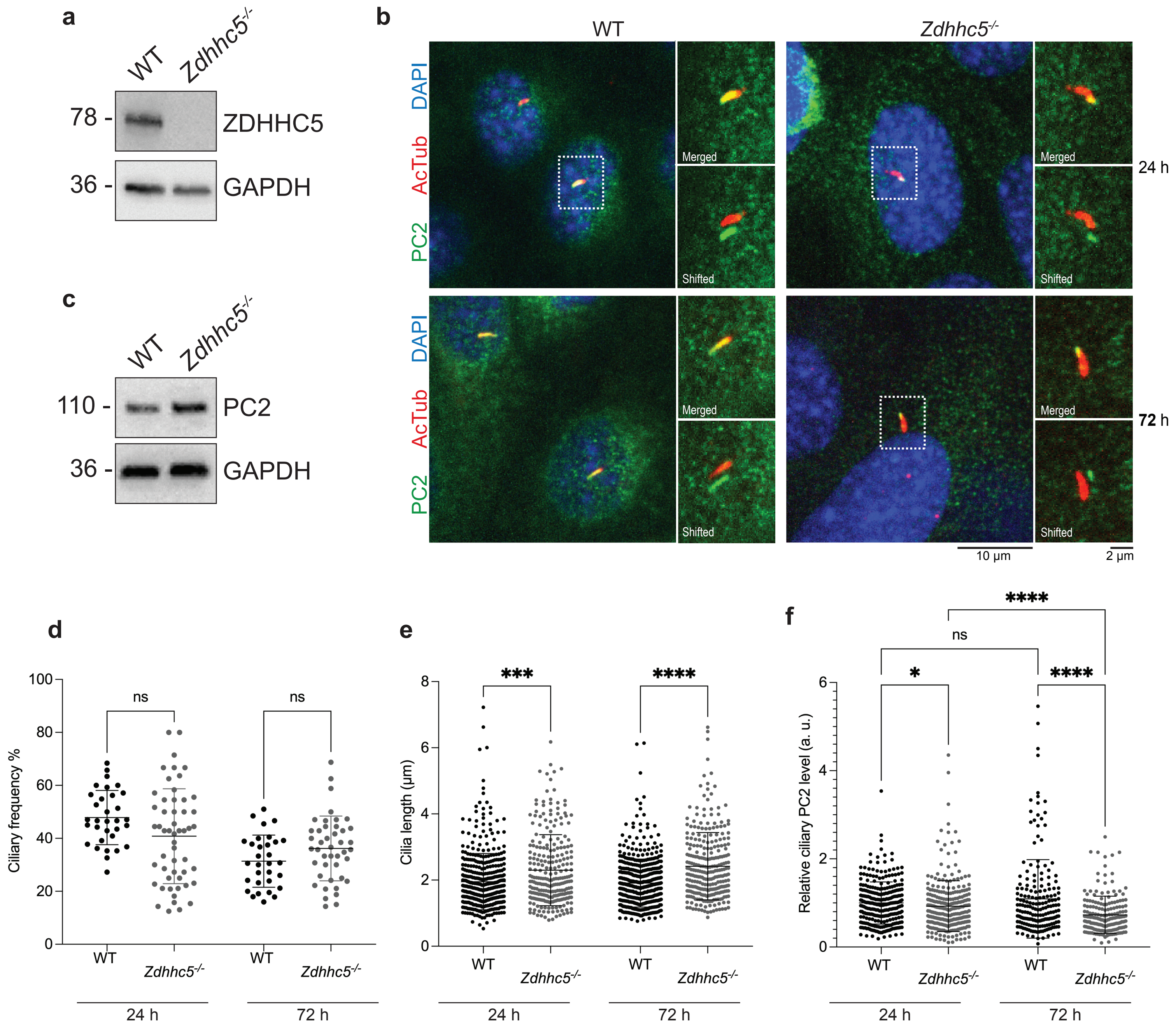
